# Flow-Induced Yap/Taz Signaling Balances Endothelial and Hematopoietic Stem Cell Fates

**DOI:** 10.64898/2026.06.20.733393

**Authors:** Wade W. Sugden, Stephan George, Zachary C. LeBlanc, Morgan T. Walcheck, Eleanor Meader, Julia Goldstein, Elizabeth Molnar, Wandi Zhu, Rubul Mout, Christopher Li, Leah Radeke, Zoey Young, Maria Gonzalez di Tillio, Marcelo Falchetti, Mohamad A. Najia, Yang Tang, Brittney D. Love, Ran Jing, Allison M. Tompkins, Olivia Stockard, Caroline Kubaczka, Katharina Kirchhof, Vanessa Lundin, Calum A. MacRae, Thorsten M. Schlaeger, George Q. Daley, Trista E. North

## Abstract

Mechanical forces from blood flow are essential for production of hematopoietic stem and progenitor cells (HSPCs) during embryogenesis, but the molecular mechanisms by which hemodynamic cues are sensed and orchestrate endothelial-to-hematopoietic (EHT) transition remain incompletely defined. We previously identified YAP mechanotransduction as a key integrator of physical forces with EHT. Here we show that hemodynamic forces can activate YAP signaling via the mechanoresponsive ion channel Piezo1 in human iPSC-derived hemogenic endothelium (HE) and zebrafish embryos. Investigation of the Piezo1/YAP axis revealed shared and unique roles of YAP and its paralogue TAZ in EHT. Mechanistically, we find a requirement for the Tead DNA-binding co-factor in YAP/TAZ-dependent control of HSPC number, and note that TAZ uniquely augments transcriptional output of the hematopoietic master regulator Runx1 via direct protein-protein interactions. By comprehensive scRNA-sequencing of YAP/TAZ gain-of-function (GOF) and yap-deficient cells from zebrafish, we reveal that YAP/TAZ promotes HSC production by positively regulating gene programs for hematopoietic self-renewal, cell cycle, and glycolysis-to-oxidative phosphorylation switching, while preventing reversion to endothelial identity. Importantly, comparison of GOF transcriptomes and functional analyses suggest decoupling of metabolic/proliferative and endothelial gene regulatory modules between YAP and TAZ: while either can functionally compensate for loss of the other in EHT, indiscriminate overactivation of TAZ enhances an endothelial program over pro-hematopoietic fate, ultimately blunting progression of HSPC production. Given that hemodynamic cues are integrated simultaneously by arterial and HE cells in embryonic vessels in which EHT occurs, these findings have strong implications for strategies designed to introduce biomechanical cues to in vitro hematopoietic differentiation systems to drive HSC production.

## Introduction

The *in vitro* production of hematopoietic stem cells (HSCs) represents a tractable target for developing cell-based therapies, based on their capability to self-renew or differentiate into progenitors capable of the production of all specialized cell types of the vertebrate blood system. Transplantation of HSCs is a curative therapy for a myriad of hematologic disorders^1–3^, and CRISPR-based gene editing of patient-derived HSCs has been demonstrated to be feasible and disease-rectifying in the clinical setting^4^. Nonetheless, despite recent advances^5,6^, a gap remains in the ability to efficiently and predictably guide directed differentiation of multilineage blood stem cells from patient-specific induced pluripotent stem cell sources (iPSC) *ex vivo* at significant scale^7^, limiting wide-spread therapeutic application. Developmentally, the potentiality and potency of the hematopoietic stem and progenitor (HSPC) pool is linked to time and tissue of origin in the embryo^8–14^, which provides strong guideposts for design of rational protocols to derive long-lived multipotent HSPCs *in vitro*.

Blood production in the embryo proceeds in successive waves. The earliest of these, the ‘primitive’ wave, produces erythroid and myeloid-competent progenitors (EMPs) to provide the earliest oxygen-transport, cytokine-directed tissue remodeling, and immune defense in the hypoxic environment of the conceptus^15^. Subsequent ‘definitive’ waves yield HSPCs that can additionally generate cells of the adaptive immune system. Definitive HSCs are endothelial in origin and have been shown to emerge from select embryonic arteries via a process termed endothelial-to-hematopoietic transition^16–18^, or EHT. Successful completion of EHT requires i) loss of the arterial endothelium gene signature^19,20^, ii) acquisition of a hematopoietic chromatin landscape and transcriptional gene program^21^, and iii) delamination (termed “budding”) from the vascular endothelial sheet and migratory capabilities to navigate to successive niches^22,23^. The gene regulatory network (GRN) responsible for the functional identity of HSCs is complex and requires a milieu of transcription factors (TFs)^24,25^; chief among these is the master regulator of hematopoietic identity Runx1^26,27^, whose expression helps demarcate hemogenic endothelium (HE) of the dorsal aorta (DA) and other embryonic arteries. The most successful iPSC to HSPC differentiation protocols to date have sought to recapitulate this developmental ontogeny to obtain mesodermal material that subsequently undergoes arterial endothelial commitment prior to HSPC emergence^6,28–32^. A recent breakthrough has succeeded in creating long-term, transplantable HSCs from iPSCs via directed differentiation of embryoid bodies in a turbulent culture system in which mesoderm patterning is guided by precise induction of retinoic acid signaling and removal of VEGF for hematopoietic progenitor expansion^5^. Gaps remain in the clonal behavior/potential of the resulting cell product, line-to-line variation in differentiation efficiency of the parent iPSCs, and the low frequency of total HSCs recovered (roughly 20-fold less than peripheral blood-derived CD34+ stem cells^5^), highlighting the need for ongoing investigation into the developmental signaling cues emerging HSCs experience during EHT to further enable translational efficacy.

Hemodynamic forces from blood flow are a critical physiologic cue sensed by endothelial cells during vascular morphogenesis^33^, which are essential for HSPC production *in vivo* in mouse^34,35^ and zebrafish embryos^36,37^, and can induce definitive hematopoiesis-associated HOX gene expression in human HE cells^38^. Key components of fluid force that impact endothelial cells are frictional wall shear stress (WSS) and orthogonal cyclic stretch (CS). Our group and others previously identified the mechanoresponsive Hippo signaling factor YAP as a key regulator of EHT^39,40^; we also demonstrated that YAP signaling is uniquely activated in HE by CS. YAP, and its paralogue WWTR1 (more commonly known as TAZ) are associated with many cellular processes in other contexts that could potentially underlie this hematopoietic fate phenotype, including cell cycling^41^, nutrient acquisition^42^, stem cell identity^43^, and migratory potential^44^. YAP/TAZ nuclear localization is tightly controlled by the phosphorylation status of conserved serine residues that either target the protein for ubiquitination/degradation or allow for nuclear import^45^. Notably, YAP/TAZ do not have direct DNA binding ability; while many of their canonical functions are ascribed to the TEAD DNA binding factor^46^, they can also interface with additional TFs, including those of the SMAD and RUNX families^47^. While YAP/RUNX2^48^ and TAZ/RUNX2^49,50^ interactions have been described in the proper differentiation of osteoblasts, the involvement of TAZ or functionality of YAP/RUNX1 or TAZ/RUNX1 interactions in hematopoietic development is unknown. Furthermore, the molecular sensors by which HE cells respond to force to initiate YAP/TAZ-associated mechanotransduction are still largely undefined.

Piezo1, a mechanically-gated ion channel, has emerged as a central force sensor in the vasculature. Endothelial deficiency of Piezo1 in mice leads to abnormal morphologic responses to hemodynamics and embryonic lethality^51^, while Piezo1 activation governs vascular responses to maintain blood pressure^52,53^. In the blood system, *PIEZO1* gain-of-function variants are associated with hereditary xerocytosis (dehydration of red blood cells), and activation of Piezo1 by the selective small molecule agonist Yoda1 slows maturation of erythroid progenitors into mature erythrocytes^54^. Recently, PIEZO1 activation was shown to promote HSPC production in mouse and human pluripotent stem cell culture^55^. Piezo1-deficient zebrafish HSPCs were notably unable to colonize the caudal hematopoietic tissue (CHT, a vascular plexus analogous to the fetal liver in mammals) effectively against wildtype (WT) competitor cells^56^. Interestingly, PIEZO1 activation has been documented to increase YAP nuclear localization and activity in a number of cell types, including osteoblasts^57^, endocardium^58^, HeLa^59^ and neuronal stem cells^60^. Whether Piezo1 can activate YAP/TAZ in HE or the mechanism by which such activity promotes EHT remain unclear.

Here, by genetic and pharmacologic manipulation, we show that hemodynamically-driven PIEZO1 activation increases both YAP activity in, and HSPC production from, human HE, and that this molecular axis is precisely conserved in zebrafish. By mapping the DNA-binding partners responsible for Yap-driven hematopoiesis, we uncover a heretofore unappreciated role for Taz, and provide evidence supporting impact of both Tead– and Runx1-directed activities of Hippo TFs in EHT. Finally, utilizing scRNA-sequencing of sorted zebrafish Kdrl+ endothelial cells from YAP loss-of-function (LOF), YAP gain-of-function (GOF), and TAZ GOF embryos, we characterize the molecular alterations in cells undergoing EHT when this mechanoregulatory axis is perturbed, revealing that hematopoietic GRN expression positively correlates with Yap dosage in HE cells. Our findings identify a conserved stretch-induced Piezo1-YAP/TAZ axis in HE, provide a mechanism by which mechanically-tuned Hippo signaling can augment lineage-specific TF function, and clarify a context-specific requirement for YAP/TAZ in specialized HE cells to maintain hematopoietic commitment during EHT in the vertebrate embryo.

## Results

### Stimulation of PIEZO1 activates YAP signaling and HSPC production in human iPSC-derived HE and zebrafish embryos

We previously showed that YAP nuclear localization could be enhanced in CD34+ human HE cells by treatment with pharmacologic Rho-GTPase activators or biomechanical stimulation, mimicking the impact of onset of blood flow in vertebrate embryos^40^. Based on observations that Piezo1 can activate the YAP axis in endothelial cells^61–63^, we hypothesized that the mechanically-gated ion channel Piezo1 may regulate YAP activity in HE. Toward this end, we treated adherent human CD34+ cells for 24hr with 0.5uM Yoda1, a small molecule Piezo1 agonist, and observed a marked increase in the nuclear-to-cytoplasmic ratio of YAP in cells exposed to Yoda1 compared to DMSO vehicle controls (**Figure 1A**, **B**). Notably, YAP expression was clearly visible in the adherent endothelial population, but absent from the floating hematopoietic cells, consistent with a role for PIEZO1 in relaying CS from blood flow in HE cells exposed to hemodynamic force. Exposure to Yoda1 increased expression of YAP target genes *ANKRD1*, *CTGF* and *CYR61* **(Figure 1C)**, while the neurotoxic peptide GsMTx-4 (a known inhibitor of Piezo1^64^) reduced expression of each in a dose-dependent manner (**Figure S1A**). Leveraging an improved *in vitro* differentiation protocol yielding highly efficient derivation of hematopoietic progenitors^31^ and our own HLF-tdTomato-reporter human iPSC line (similar to existing constructs^65^) (CL, MN, GQD; *unpublished*), we assessed the effect of stimulating Piezo1 in CD34+ HE on yield of functional hematopoietic progenitors. Treatment with Yoda1 led to increased numbers of cells expressing the HLF-tdTomato HSC reporter and the fraction of CD34+/CD45+ hematopoietic cells in the cultures (**Figure 1D-F** and **S1B, C**).

**Figure 1:**
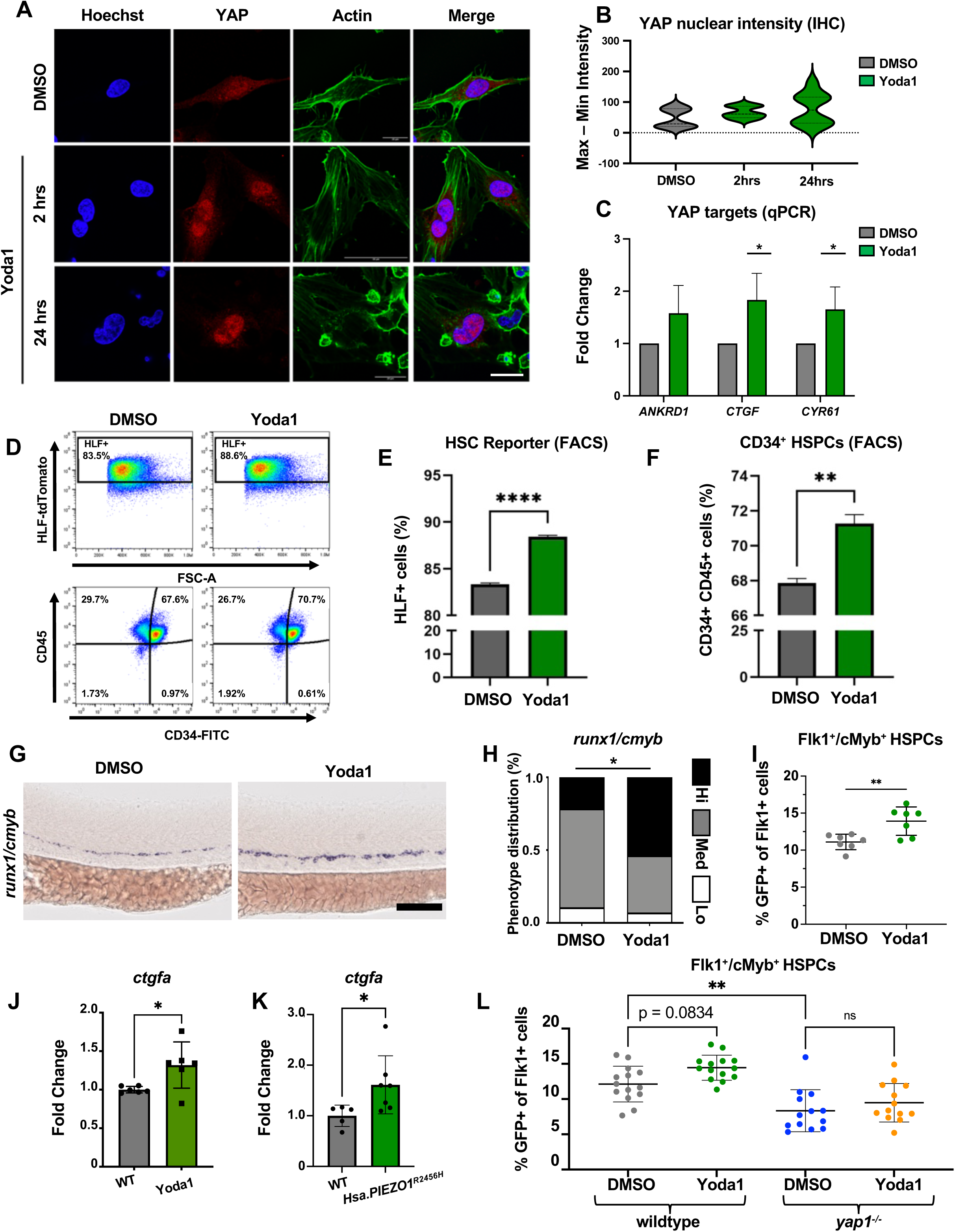
Piezo1-YAP axis can be pharmacologically stimulated to drive HSPC production in human HE cells and zebrafish. **A**) Maximum intensity projection of confocal z-stack of immunohistochemistry for Hoechst (*blue*), YAP (*red*), and F-actin (*green*) in human 2D-derived CD34+ hemogenic endothelial cells exposed to DMSO or the Piezo1 agonist Yoda1 (0.5uM) for 2hrs and 24hrs. Scale bar: 20um. **B)** Quantification of nuclear/cytoplasmic ratio of YAP staining from images in (**A**) (n ≥ 8). **C)** qPCR for human YAP target genes in human CD34+ cells exposed to DMSO or Yoda1 for 24hrs. Expression normalized to *GAPDH* housekeeping gene (n = 3; 1-way ANOVA, *p ≤ 0.05). **D)** Representative flow plots of HLF-tdTomato reporter and CD34/CD45 surface marker (antibody) expression in hemogenic endothelial cells exposed to DMSO or Yoda1 (0.5uM) from Day 5-7 of differentiation. **E, F)** Quantification of fraction of human iPSC-derived cells expressing HLF reporter (n = 3, unpaired Student’s t-test, ****p ≤ 0.001) and CD34+/45+ (n=3; unpaired Student’s t-test, **p ≤ 0.01) following DMSO or Yoda1 treatment as in (**D**). **G, H)** *runx1/cmyb* WISH and qualitative distribution plot in WT embryos exposed to DMSO or Yoda1 (1uM) from 10 somite stage (ss) –36hpf and scored with low (lo), medium (med), or high (hi) *runx1* expression in the VDA (n ≥ 25 embryos/condition; chi-squared test, *p ≤ 0.05). Scale bar: 100um. **I)** FACS analysis of Flk1+/cMyb+ HSPCs in 48hpf *Tg*(*kdrl*:*Hsa*.*HRAS-mCherry*)*^s916^*; *Tg*(*cmyb*: *EGFP*)*^zf169^* reporter embryos exposed to Yoda1 (0.5uM) from 24-48hpf (n = 7 pools of embryos/condition, 7 clutches); unpaired Student’s t-test, *** p ≤ 0.001; Error bars indicate SD). **J)** Whole-embryo qPCR for zebrafish *ctgfa* in WT embryos exposed to DMSO or Yoda1 (2uM) from 10ss-36hpf. Expression normalized to *18S* housekeeping gene (n = 6; unpaired Student’s t-test, *p ≤ 0.05; Error bars indicate SD). **K)** Whole-embryo qPCR for zebrafish *ctgfa* in WT and *Tg(ef1a:Hsa.PIEZO1^R2456H^)* embryos at 5dpf. Expression normalized to *eef1a* housekeeping gene (n = 4-7; unpaired Student’s t-test, *p ≤ 0.05; Error bars indicate SD). **L)** FACS analysis of Flk1+/cMyb+ HSPCs in 48hpf transgenic WT and *yap1^−/−^* embryos exposed to DMSO or Yoda1 (0.5uM) from 24-48hpf (n = 14; one-way ANOVA, *p ≤ 0.05, ***p ≤ 0.001; Error bars indicate SD).

To determine if this Piezo1/YAP axis was conserved and could regulate HSPC numbers *in vivo*, we employed the genetically tractable zebrafish system. Zebrafish embryos treated from 14-36 hours post fertilization (hpf) with 1uM Yoda1 showed increased expression of the conserved HSPC markers *runx1* and *cmyb* in the DA by combinatorial whole-mount in situ hybridization (WISH) (**Figure 1G, H**). A positive impact on *runx1/cmyb* expression was evident across a dose curve of Yoda1 exposure (**Figure S1D**). This finding was validated by FACS, where Yoda1 treatment (0.5uM, 24-48hpf) significantly increased Flk+/cMyb+ HSPCs in *Tg*(*kdrl*:*Hsa*.*HRAS-mCherry*)*^s916^*;*Tg*(*cmyb*:*EGFP*)*^zf169^*reporter embryos (**Figure 1I** and **S1E**). Notably, upregulation of the YAP target gene *ctgfa* was observed following Yoda1 treatment as well as in embryos globally overexpressing a human gain-of-function (GOF) PIEZO variant (**Figure 1J, K**). Epistasis experiments indicated that while Yoda1 increased the number of Flk+/cMyb+ HSPCs in wildtype (WT) embryos, Yoda1 treatment could not rescue the significant deficiency in HSPC number found in *yap1^mw48^* mutant embryos (hereafter referred to as *yap1^−/−^*) (**Figure 1L**). Collectively, these results identify PIEZO1 as a conserved cellular mediator that can stimulate the production of HSPCs in human cells and in zebrafish embryos, at least in part, by activation of YAP.

### Piezo1 is a sensor of cyclic stretch to drive YAP/TAZ-dependent HSPC production *in vivo*

We next asked if Piezo1/Yap regulates hematopoietic development downstream of blood flow *in vivo*. Double fluorescent WISH confirmed that *piezo1* is expressed in the DA at ∼32hpf (during the developmental window of EHT), including in *runx1*+ HE cells (**Figure S2A**). Surprisingly, analysis of *piezo1^um136^* mutant embryos (hereafter referred to as *piezo1^−/−^*) derived from *piezo1^+/−^*incrosses initially revealed no phenotype: embryos were indistinguishable from WT siblings, with no obvious defects in circulation or hemodynamics, and no change in *runx1/cmyb* expression at 36hpf (**Figure 2A, top**). To uncouple the contribution of WSS and CS forces in the DA, we employed a pharmacological “stretch only” paradigm whereby embryos were titrated with pimozide, a dopamine receptor antagonist known to impede cardiac output^66^, to a dose that visibly eliminated erythrocyte circulation (a proxy for WSS) but did not completely abolish irregular heart contractions (**Figure S2B, C**). Embryos experiencing this ‘stretch only’ profile maintained *runx1/cmyb* expression closer to WT levels than that seen in *tnnt2a silent heart* morpholino (MO) injected embryos^67^, which lack all hemodynamic cues (**Figure S2D).** This phenotype is similar to published observations of *malbec/cdh5* mutants that maintain normal HE specification despite the lack of circulatory flow by virtue of residual heartbeats that propagate stretch forces through the plasma^68^. Significantly, *piezo1^−/−^* embryos were sensitized to this experimental ‘stretch only’ manipulation and exhibited reduced *runx1/cmyb* staining compared to WT controls (**Figure 2A, bottom**).

**Figure 2:**
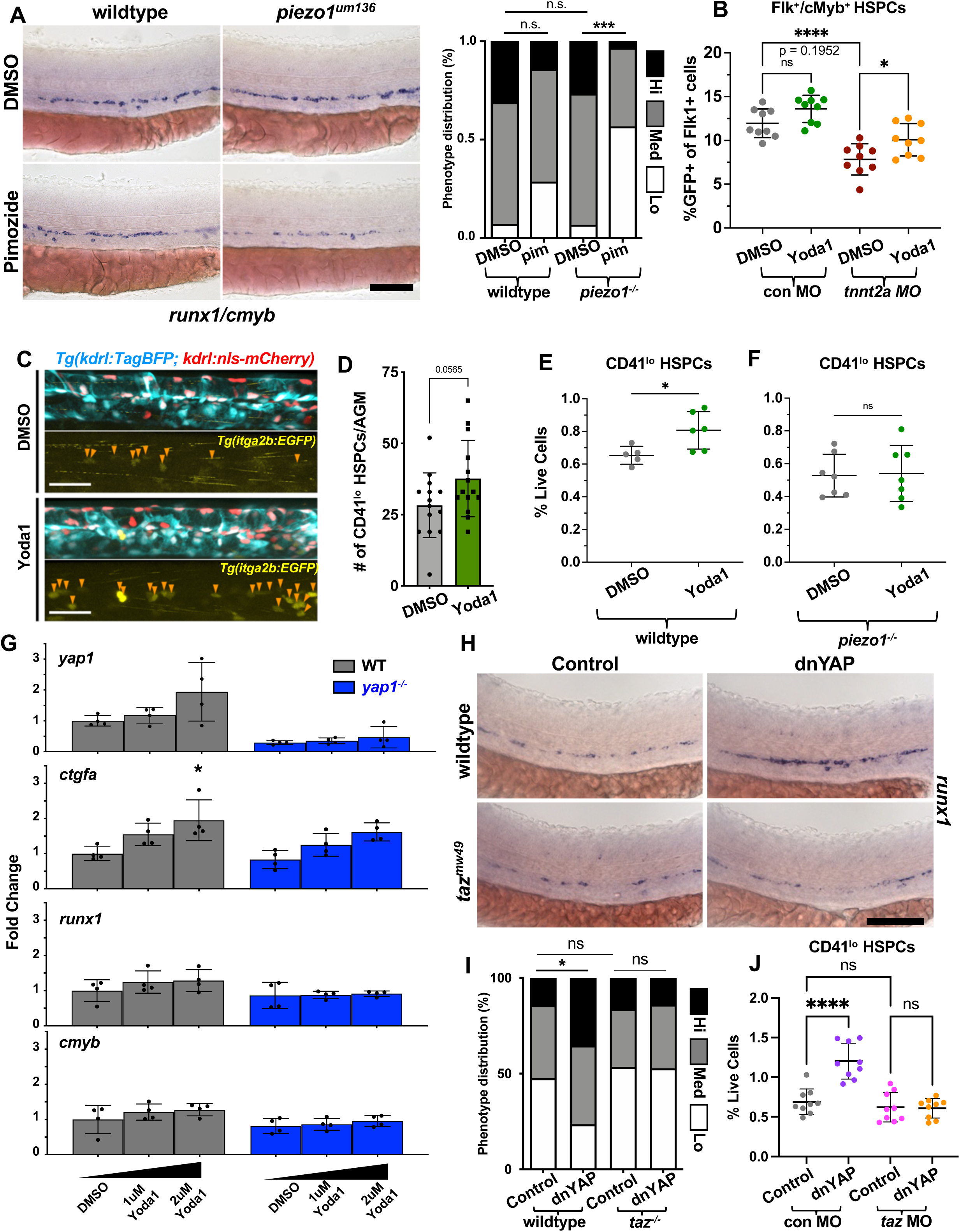
Piezo1 relays cue from hemodynamic cyclic stretch to YAP/TAZ signaling. **A**) Representative *runx1/cmyb* WISH and qualitative distribution plot in WT and *piezo1^−/−^* embryos exposed to DMSO or 25uM pimozide from 24-36hpf and scored with low (lo), medium (med), or high (hi) *runx1* expression in the VDA (n ≥ 28 embryos/condition; chi-squared test, n.s. = not significant, ***p ≤ 0.001). Scale bar: 100um. **B)** FACS analysis of Flk1+/cMyb+ HSPCs in 48hpf standard control and *silent heart/tnnt2a* MO embryos exposed to DMSO or Yoda1 (0.5uM) (n = 9; one-way ANOVA, n.s. = not significant, *p ≤ 0.05, **** p ≤ 0.0001; Error bars indicate SD). **C)** Representative maximum intensity projection of confocal z-stack of CD41:EGFP+ HSPCs (yellow) in dorsal aorta at 48hpf in DMSO or Yoda1 (2uM) treated embryos (dosed from 10ss-48hpf). Endothelial cytoplasm and nuclei labeled by *Tg(kdrl:TagBFP)^mu293^* (cyan) and *Tg(kdrl:nls-mCherry)^is4^*(red) transgenes, respectively. Scale bar: 50um **D)** Quantification of CD41:EGFP+ cells in the VDA from embryos treated as in C (n = 14; unpaired Student’s t-test; Error bars indicate SD). **E, F)** FACS analysis of CD41:EGFP^lo^ HSPCs in 72hpf transgenic WT and *piezo1^−/−^* embryos exposed to DMSO or Yoda1 (0.5uM) (n = 6 WT, 7 mutant; unpaired Student’s t-test, *p ≤ 0.05; Error bars indicate SD). **G)** Whole-embryo qPCR on RNA from WT and *yap1^−/−^* embryos at 30hpf. Dose curve of Yoda1 treatment was applied from 10ss–30hpf. YAP target gene *ctgfa* expression increases in stepwise fashion, with similar trends for *yap1, runx1* and *cmyb.* Expression normalized to *18S* housekeeping gene (n = 4; one-way ANOVA, *p ≤ 0.05; Error bars indicate SD). **H)** WISH for *runx1* at 30hpf in WT and *taz^−/−^* embryos with or without HS:dnYAP transgene induction by heat-shock (HS) at 24hpf. Scale bar: 100um. **I)** Qualitative phenotypic distribution plot of embryos in (I) scored with lo/med/hi *runx1* expression in the VDA at 30 hpf (n = 42 WT control, 43 *taz^−/−^*, 34 dnYAP, 36 *taz^−/−^*/dnYAP; chi-squared test, n.s. = not significant, *p ≤ 0.05). **J)** FACS analysis of CD41:EGFP^lo^ HSPCs at 72hpf in control and dnYAP embryos injected with standard control or *taz* MO, with HS administered at 24hpf (n = 9; unpaired Student’s t test, n.s. = not significant, ****p ≤ 0.0001; Error bars indicate SD).

Consistent with Piezo1 functioning downstream of the mechanical input from blood flow to drive HSPC production from HE, we observed that treatment of *tnnt2a* morphants with Yoda1 to directly activate Piezo1, led to a partial rescue of Flk+/cMyb+ HSPCs (**Figure 2B**). Confocal imaging of WT embryos exposed to Yoda1 from the 10 somite stage (ss) to 48hpf confirmed an increased proportion of EGFP^lo^ budding EHT cells in the floor of the DA of transgenic *Tg(–6.0itga2b:EGFP)^la2^* [referred to as CD41:EGFP] reporter embryos (**Figure 2C, D**). FACS analysis of CD41^lo^ HSPCs at 72hpf revealed that the Yoda1-driven increase in total HSPCs was Piezo1-dependent, confirming both compound specificity and the loss-of-function nature of the *um136* allele (**Figure 2E, F** and **S2E**). These results indicate that Piezo1 functions as an active membrane mechanosensor regulating HSPC development by sensing vascular CS, and that this function is likely augmented by additional mechanosensors and/or compensatory contributions mediated by WSS in the DA.

Having established that Piezo1 could function as a mechanosensor *in vivo*, we returned to the question of how it might promote HSPC production via YAP activation. WT embryos displayed a dose-dependent increase in the YAP target gene *ctgfa* and hematopoietic transcription factors *runx1* and *cmyb* in response to Yoda1 at 30hpf; these changes were notably abrogated in *yap1^−/−^* embryos, with the exception of *ctgfa* expression, which continued to increase (**Figure 2G**). We suspected this outcome might be due to Piezo1-driven stimulation of TAZ-associated mechanotransduction: TAZ is a paralogue of YAP, and the two proteins have a suite of redundant and unique transcriptional outputs depending on cell-type^69^. To circumvent gross developmental defects caused by compound *yap/taz* genetic deficiency during early development^70^, we first characterized hematopoietic development in embryos expressing a heat shock-inducible dominant negative YAP (hereafter referred to as dnYAP) as a means to temporally modulate YAP activity. This variant contains a TEAD binding domain (TBD) and two WW domains of zebrafish YAP1 protein fused to a nuclear localization signal (NLS), but lacks the transactivation domain (**Figure S2F**). The resulting protein is predicted to localize to the nucleus and interact with YAP DNA-binding partners, but not regulate transcription; it had been previously documented to recapitulate YAP LOF phenotypes in chicken models of neurogenesis^71^. Unexpectedly, induction of dnYAP at 10ss led to increased expression of HSPC markers *runx1/cmyb* at 30hpf by WISH (**Figure S2G, H**). Furthermore, heat shock induction at 24hpf produced a quantifiable increase in CD41:EGFP^lo^ HSPCs by FACS at 72hpf (**Figure S2I**).We hypothesized that the ‘hematopoietic rebound’ effect observed in dnYAP embryos might be due to a latent role for TAZ. In line with our previous findings^40^, whole embryo qPCR analysis revealed an increase in canonical YAP/TAZ target genes *ctgfa* and *amotl2b* following overexpression of a heat shock-inducible constitutively active variant of zebrafish Yap1 (YAP1^S87A^, hereafter referred to as caYAP), and indicated that dnYAP also upregulated these genes about half as efficiently (**Figure S2J**). Therefore, we next evaluated HE specification and HSPC production in *taz* loss-of-function animals in the context of dnYAP induction. We performed dnYAP overexpression at 24hpf in embryos from incrosses of *taz^mw49+/−^*adults, followed by *runx1* ISH at 30hpf and FACS analysis of CD41^lo^ HSPCs at 72hpf. Strikingly, while *runx1* expression in *taz^−/−^* embryos was indistinguishable from controls at baseline, loss of TAZ protein completely abrogated the increased *runx1* expression found in dnYAP embryos (**Figure 2H, I**). We saw the same Taz dependency for the dnYAP-driven increase in CD41^lo^ HSPCs at 72hpf, which could be curbed by administration of 1ng *taz* MO (**Figure 2J**)). Higher doses of *taz* MO did decrease HSPC numbers compared to control injections (**Figure S2K**), leaving open the possibility that redundant Yap/Taz activity in HE is masked by genetic compensation^72^ when either one of the two TFs is lost. Together, these data provide the first *in vivo* evidence for a specific role of the Taz transcription factor in definitive hematopoiesis, and emphasize a critical window of function for Yap/Taz regulation during the window of hemodynamic flow through the DA, which can be sensed by Piezo1, to promote HSPC formation.

### Yap and Taz control HSPC production via Tead-directed and Runx1-directed regulatory activity

As Yap/Taz function as transcriptional cofactors, we next wished to determine which DNA binding proteins are responsible for Yap/Taz-dependent hematopoiesis. YAP and TAZ share highly similar protein structure: both initiate with a TBD, followed by one or two WW domains and a transactivation domain (**Figure 3A**). Eight transcriptional isoforms of YAP have been described in human cells, with the nomenclature YAP1-1 and YAP1-2 used to refer to the number of WW domains encoded in the transcript^73^. In contrast, TAZ has a single transcriptional isoform containing one WW domain, most similar to the first WW domain of YAP^74^. Given the critical importance of RUNX1 as a master regulator of hematopoietic fate in HE, and the reported regulatory interaction of the RUNX family with YAP/TAZ^47–50^, we set out to evaluate the requirement for TEAD and RUNX1 in YAP/TAZ-dependent transcription, using human YAP/TAZ variants with diminished ability to interact with select DNA binding partners (**Figure 3B**). Utilizing a well-characterized 8xGTIIC-LUC construct driven by TEAD enhancers^75^, we observed strong induction of luciferase activity by both human YAP and TAZ in HEK293 cells (**Figure 3C**). Importantly, this induction was refractory to mutations in the WW domains of YAP, but abolished by mutations that disrupt the YAP/TEAD interface, confirming specificity of the assay. Next, we asked if either YAP or TAZ, when co-expressed with RUNX1, could augment transcription from RUNX1 enhancers using a 6xOSE-LUC construct (originally derived from RUNX2 enhancers, sharing the common DNA binding motif for RUNX proteins)^76^. Surprisingly, we observed synergistic regulation from RUNX1/TAZ, but not RUNX1/YAP, in promoting RUNX1-dependent expression from 6xOSE-LUC (**Figure 3D**). Given the high degree of sequence and structural similarity between YAP and TAZ, we aimed to further investigate the specificity of this relationship. We confirmed that, *in vitro*, YAP1-1 and TAZ proteins could drive robust TEAD-dependent transcription (YAP1-1 less efficiently than TAZ), and that mutations in the WW domain of TAZ did not significantly impact TAZ/TEAD signaling strength (**Figure 3E**). Strikingly however, we saw no enhancement of Runx1-dependent luciferase signal when co-expressing RUNX1 and YAP1-1 (**Figure 3F**), despite this variant having almost 52% identical amino acid homology (∼70% similarity) to TAZ. Furthermore, this synergistic increase was blunted by introduction of mutations predicted to interfere with the interface between the TAZ WW domain and PPXY motif in RUNX1. Together, these findings underscore the uniqueness of RUNX1/YAP versus RUNX1/TAZ functional unit particularly on transactivation of RUNX enhancers.

**Figure 3:**
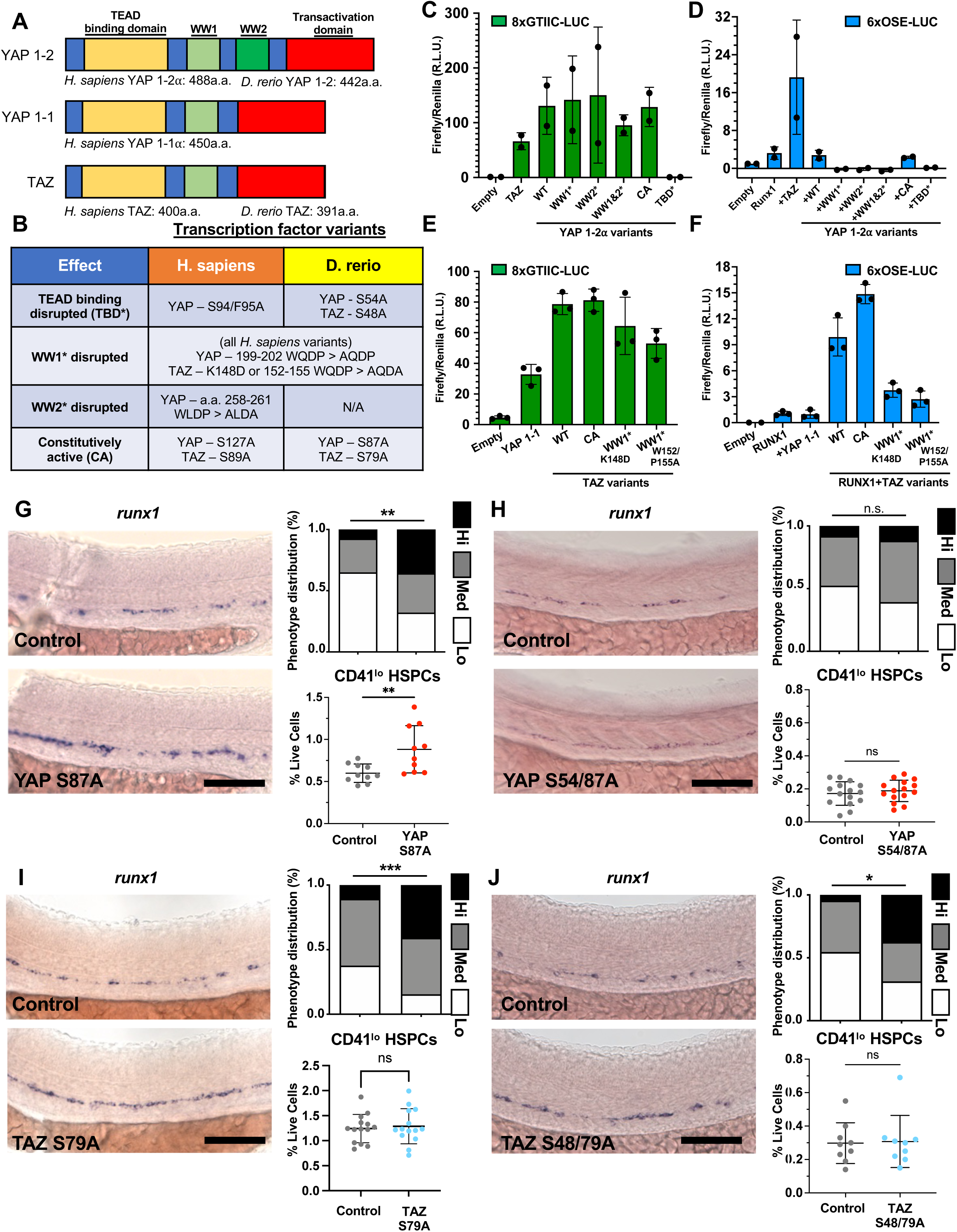
Functional experiments support TEAD– and Runx1-driven YAP/TAZ activity in hematopoiesis. **A, B**) Schematic of YAP/TAZ isoform protein structure and summary of variants used in this study. **C)** Activity of human YAP/TAZ variants on TEAD-driven 8xGTIIC-LUC reporter in HEK293T cells (n = 2; Error bars indicate SD). **D)** Activity of human RUNX1 alone or with human YAP/TAZ variants on RUNX-driven 6xOSE-LUC reporter in HEK293T cells (n = 2; Error bars indicate SD). **E)** Activity of human single WW domain YAP/TAZ variants on TEAD-driven 8xGTIIC-LUC reporter in HEK293T cells (n = 3; Error bars indicate SD). **F)** Activity of human single WW domain YAP/TAZ variants on RUNX1-driven 6xOSE-LUC reporter in HEK293T cells (n = 3; Error bars indicate SD). **G)** WISH for *runx1* expression (*left*) with qualitative phenotypic distribution plot at 30hpf (*upper right*), and FACS analysis of CD41:EGFP^lo^ HSPCs (*lower right*) at 72hpf in control and HS:YAPS87A embryos (*constitutively active*). HS at 24hpf (for WISH: n ≥ 40 embryos/condition; chi-squared test, **p ≤ 0.01; for FACS: n = 10; unpaired Student’s t-test, **p ≤ 0.01; Error bars indicate SD). Scale bar: 100um. **H)** WISH for *runx1* expression (*left*) with qualitative phenotypic distribution plot at 30hpf (*upper right)*, and FACS analysis of CD41:EGFP^lo^ HSPCs (*lower right*) at 72hpf in control and HS:YAPS54/87A embryos (*cannot bind TEAD*). HS at 24hpf (for WISH: n ≥ 60 embryos/condition; chi-squared test, n.s. = not significant; for FACS: n = 15; unpaired Student’s t-test, n.s. = not significant; Error bars indicate SD). Scale bar: 100um. **I)** WISH for *runx1* expression (*left*) with qualitative phenotypic distribution plot at 30hpf (*upper right)*, and FACS analysis of CD41:EGFP^lo^ HSPCs (*lower right*) at 72hpf in control and HS:TAZS79A embryos (constitutively active). HS administered at 24hpf (for WISH: n ≥ 60 embryos/condition; chi-squared test ***p ≤ 0.001; for FACS: n = 8; unpaired Student’s t-test **p ≤ 0.01; Error bars indicate SD). Scale bar: 100um. **J)** WISH for *runx1* expression (*left*) with qualitative phenotypic distribution plot at 30hpf (*upper right)*, and FACS analysis of CD41:EGFP^lo^ HSPCs (*lower right*) at 72hpf in control and HS:TAZS48/79A embryos (*cannot bind TEAD*). HS administered at 24hpf (for WISH: n ≥ 16 embryos/condition; chi-squared test, *p ≤ 0.05; for FACS: n = 9; unpaired Student’s t-test, n.s. = not significant; Error bars indicate SD). Scale bar: 100um.

We then turned back to the zebrafish model to assess how modulating the TEAD and/or RUNX1 axis would impact the effect of YAP and/or TAZ on hematopoietic development *in vivo*. First, we generated a heat-shock inducible constitutively active variant of zebrafish TAZ (hereafter referred to as caTAZ), and confirmed its ability to upregulate YAP/TAZ target genes by whole-embryo qPCR (**Figure S3A**). Next, we created additional variants of caYAP and caTAZ that disrupt their ability to interact with Tead (S54 and S48, respectively), and compared how they impacted YAP/TAZ GOF effects on hematopoietic specification (*runx1* expression at 30hpf by WISH) and HSPC numbers (FACS for CD41:EGFP^lo^ HSPCs at 72hpf). All variants were overexpressed at 24hpf. Induction of caYAP increased both *runx1+* HE cells at 24hpf and phenotypic HSPCs at 72hpf (**Figure 3G**); neither of these phenotypes were observed with YAP S54/87A overexpression (**Figure 3H**), indicating dependence on TEAD binding. Conversely, caTAZ overexpression led to increased *runx1* levels at 30hpf, with no change in HSPC numbers at 72hpf (**Figure 3I**). Furthermore, increased expression of *runx1* at 30hpf was still observed in TAZ S48/79A embryos, which feature disruption of the TEAD-binding interface, with no impact on HSPC numbers at 72hpf (**Figure 3J**). Interestingly, caTAZ overexpression restored HSPCs in *yap1* MO embryos to WT levels (**Figure S3B**), and synthetically tethering YAP or TAZ to RUNX1-bound loci by inducing Runx1^dbd^-Yap or Runx1^dbd^-Taz fusion proteins noticeably increased *cmyb* expression and HSPC production in the embryo **(Figure S3C, D)**. Taken together, these data offer insight into complex requirements for both YAP and TAZ in regulating EHT. Overlapping functions allow one TF to compensate for total loss of the other, but in the context of overexpression, unique transcriptional outputs have divergent impacts on HSPC production. Our analysis of YAP/TAZ variants supports a role for both TEAD– and RUNX1-directed activities of YAP and TAZ in hematopoietic development, and suggest that the ability of RUNX1/TAZ to regulate HE specification is uncoupled from TEAD-driven effects on phenotypic HSPC number.

### Yap augments expression of hematopoietic self-renewal gene regulatory network in hemogenic endothelial cells

To understand the molecular basis for the differential effects of YAP and TAZ on EHT and HSPC production, we employed a scRNA-sequencing approach to profile the transcriptional landscape of Kdrl+ endothelial-derived cells collected from *yap1^−/−^* loss-of-function (LOF) embryos and GOF embryos overexpressing caYAP. (**Figure 4A**). Cells were sorted from dissected and disaggregated zebrafish tails from transgenic *kdrl* reporter embryos for each genotype (and corresponding stage-matched WT controls) at ∼36hpf, enriching for the anatomical location of HSPC emergence at the peak of budding from the DA. Clustering yielded 24 populations, including mesodermally-derived blood and vascular cells, as well as several non-vascular populations with epidermal, neural and sclerotome signatures, consistent with previous reports^77^ (**Figure S4A, B**). We subselected hematovascular populations, inclusive of venous endothelial cells (VECs), arterial endothelial cells (AECs), hematopoietic stem/progenitor cells (HSPCs), erythroid– (E-HSPCs) and myeloid-(M-HSPCs) progenitors (similar to published datasets^78^), committed erythromyeloid cells (EMCs), and mature erythroid cell clusters for further analysis (**Figure 4B**).

**Figure 4:**
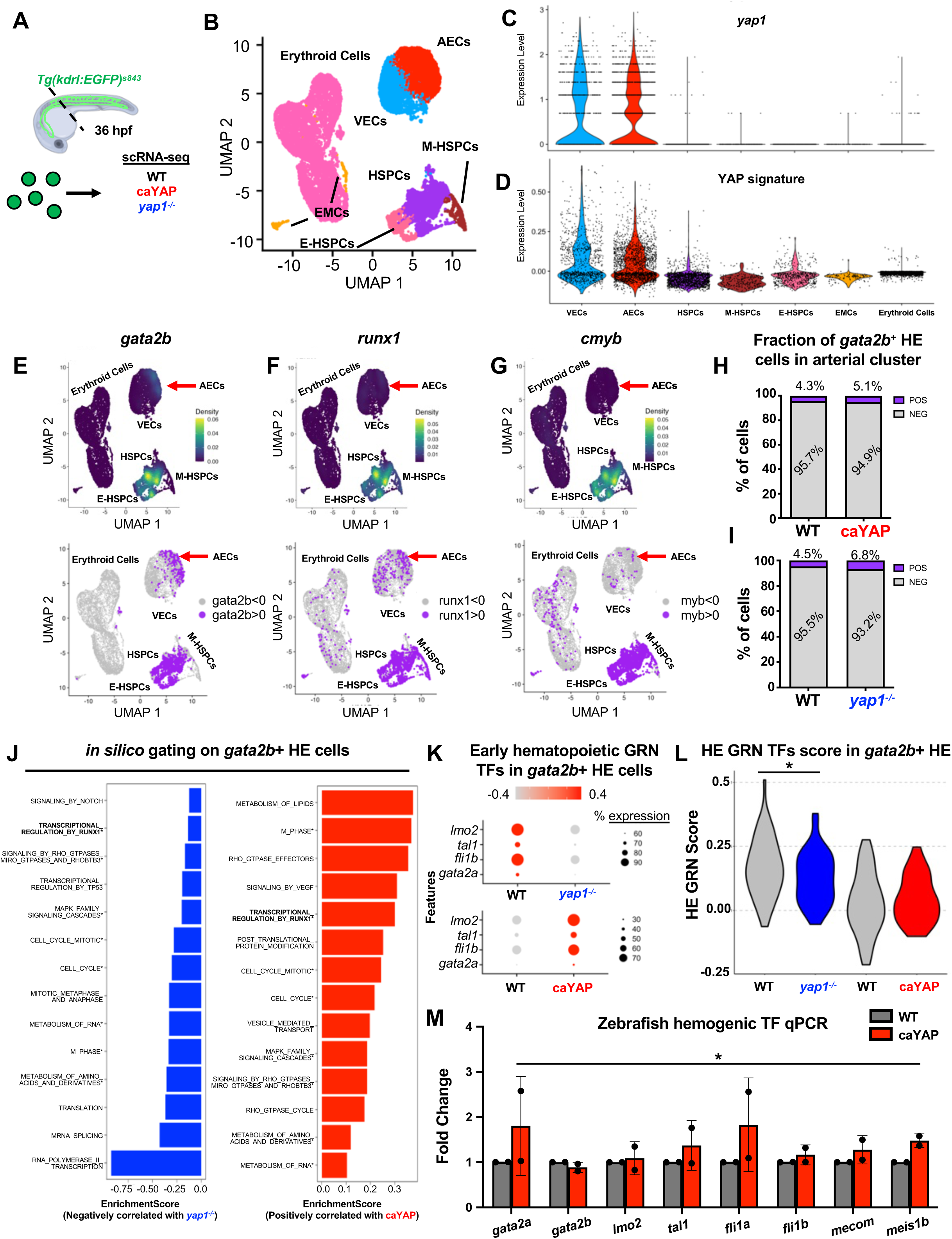
YAP promotes self-renewal and stemness in early hemogenic endothelium. **A**) Sorting strategy for scRNA-seq profiling of WT, YAPS87A (constitutively active YAP, caYAP) and *yap1^−/−^*mutant Kdrl+ endothelial cells from dissected trunks at 36hpf. **B)** Combined integrated UMAP of reclustered Kdrl+ hematovascular populations from GOF and LOF experiments. Abbreviations: VEC=Venous Endothelial Cell, AEC=Arterial Endothelial Cell, HSPCs=Hematopoietic Stem and Progenitor Cells, M-HSPCs=Myeloid-biased HSPCs, E-HSPCs=Erythroid-biased HSPCs, EMCs=Erythromyeloid Cells. **C)** Violin plots of *yap1* expression across endothelial and hematopoietic UMAP clusters. **D)** Violin plots YAP signaling ‘score’ from expression module of 9 target genes across endothelial and hematopoietic UMAP clusters. **E-G)** Density plots showing expression of hematopoietic transcription factors *gata2b, runx1* and *cmyb* (top), with cells defined as positive/negative by threshold (bottom). All TFs are highly expressed in the HSPC cluster, but *gata2b* has a noticeable distribution in a subpopulation of the arterial cells (red arrow), the presumptive hemogenic endothelium (HE). **H)** Quantification of *gata2b+* and *gata2b-* cells within the arterial cluster for WT and caYAP samples in the GOF experiments (n = 883 total WT cells, 704 total caYAP cells). **I)** Quantification of *gata2b+* and *gata2b-* cells within the arterial cluster for WT and *yap1^−/−^* samples in the LOF experiments (n = 1156 total WT cells, 818 total yap1^−/−^ cells). **J)** GSEA analysis for Reactome pathways enriched among differentially expressed genes in *gata2b+* HE cells from WT vs *yap1^−/−^* mutant and WT vs caYAP samples. The terms ‘Transcriptional_Regulation_by_Runx1’ and Cell Cycling are associated with the genes positively regulated by the YAP transcription factor (TF). **K)** Dot plot of hematopoietic triad and early hematopoietic TF expression in *gata2b+* HE cells from WT vs *yap1^−/−^*and WT vs caYAP samples. All genes are upregulated by caYAP overexpression, and most are reduced in *yap1^−/−^* cells. **L)** Violin plot of composite hematopoietic GRN ‘score’ in *gata2b+* HE cells in GOF and LOF experiments. Derived from a module of 8 hematopoietic TFs enriched in HSCs and involved in self-renewal, this score positively correlates with YAP dosage (n = 54 WT, 60 *yap1^−/−^* cells for LOF, 39 WT, 37 caYAP cells for GOF; unpaired Student’s t-test, *p ≤ 0.05). **M)** Whole-embryo qPCR for a panel of hematopoietic TFs at 30hpf in WT and caYAP embryos following HS at 24hpf. Expression normalized to *18S* housekeeping gene (n = 2; full panel evaluated by two-way ANOVA, *p<0.05; Error bars indicate SD).

Exploration of the WT cells mirrored known expression patterns and Yap activity: Yap expression was highest in VECs and AECs, and largely undetectable in HSPCs (**Figure 4C**). We saw a concomitant reduction in the strength of expression of ‘Yap signature’ genes across VEC-AEC-HSPC-erythroid clusters (**Figure 4D**). Analysis of the master hematopoietic regulator TFs *gata2b, runx1*, and *cmyb* showed high expression levels of each in the HSPC cluster (**Figure 4E-G)**. Notably, *gata2b* was unique in having a sizeable additional distribution within the AEC cluster (**Figure 4E**, red arrow), consistent with its ability to demarcate the pre-hemogenic endothelium^79^. Yap dosage did not drastically alter the percentage of AEC cells expressing *gata2b,* with approximately 4-5% of AEC cells being defined as *gata2b*+ with expression level >3 in the integrated data (**Figure 4H**). A slight increase in this fraction was noted in *yap1^−/−^* embryos, which could ostensibly be due to a block in further differentiation (**Figure 4I**) or a shift to erythroid fate^80^. By setting an *in silico* gate on these presumptive HE cells and performing differential expression analysis, we observed inverse gene expression changes in the LOF and GOF samples that converged on similar biological processes. GSEA analysis against the Reactome database^81^ for genes that positively correlated with Yap dosage revealed overlapping terms related to cell cycle and metabolism. Most strikingly, the term “transcriptional regulation by Runx1” was inversely shared by the GOF and LOF samples (**Figure 4J**), underscoring a potential context-specific function for Yap in augmenting the impact of Runx1 transcriptional function in HE.

Runx1 is essential for de novo HSC formation via EHT but considered largely dispensable afterwards^82^. It is known to form a key node in the hematopoietic GRN that secures hematopoietic fate in HE cells^83^. To that end, we assessed a curated list of TFs known to be upstream of RUNX1 in forming this GRN, many of which are coordinately regulated by RUNX1 itself, as well as TFs associated with HSC emergence. We saw increased expression of the hematopoietic ‘triad’ of HSC commitment TFs: *gata2*, *fli1* and *scl/tal1*^84^, as well as *lmo2* in HE overexpressing caYAP; the majority of which were conversely downregulated by loss of Yap (**Figure 4K**). We created a hemogenic GRN module score by combining *lmo2, tal1, fli1a, fli1b, gata2a* and *gata2b* with hemogenic transcription factors *meis1b* and *mecom*^85^, revealing a positive correlation between YAP dosage and HE commitment (**Figure 4L**). Strikingly, evaluation of genes associated with more differentiated hematopoietic cell types showed the opposite trend: erythroid genes, in particular, were generally suppressed by caYAP in HE (**Figure S4C**), while an erythromyeloid signature was elevated in *yap1^−/−^*cells (**Figure S4D**). These effects were validated by qPCR, whereby YAP overexpression upregulated a battery of TFs essential to promote the hematopoietic self-renewal GRN at the whole-embryo level (**Figure 4M**). Notably, this effect was pharmacologically recapitulated by Piezo1 activation via Yoda1 treatment (24-30hpf) in zebrafish embryos (RNA measured from dissected tails) (**Figure S4E**). Yoda1 treatment also upregulated the hematopoietic TFs *RUNX1* and *GFI1B* in human iPSC-derived CD34+ HE cells (**Figure S4F**). Together, these data suggest that YAP functions in a context-specific manner in HE to promote the expression of hematopoietic GRN TFs, including augmenting the RUNX1 regulatory signature, leading to enhancement of a ‘stemness’ state while preventing premature differentiation.

### Yap deficiency in HE causes reversion to endothelial identity and failure of glycolysis-to-oxidative phosphorylation metabolic transition

Previous work suggested that loss of hemodynamics in mouse embryos causes a differentiation block in EHT^35^. To determine how loss of Yap function *in vivo* might affect hematopoietic progression, we performed supervised trajectory analysis with Tradeseq^86^ to array the cells along developmental pseudotime recapitulating the biological ontogeny of HSPCs, descending from endothelial precursors, with erythroid cells included as a representative terminal state (**Figure 5A**). Notably, we observed a shift toward fewer *yap1^−/−^* cells in the HSPC peak of the trajectory, whereas there was a slight increase in mutant cells in the erythroid fraction (**Figure 5B**). Visual inspection of *gata1a* expression revealed that *yap1^−/−^* mutants show notably higher expression of this erythroid lineage master regulator in the AEC population than WT (**Figure S5A**); similar results were observed when plotting pseudotime trajectories with monocle (**Figure S5B**). These findings indicate that erythroid lineage fate, in particular, is normally restrained in HE by Yap activity.

**Figure 5:**
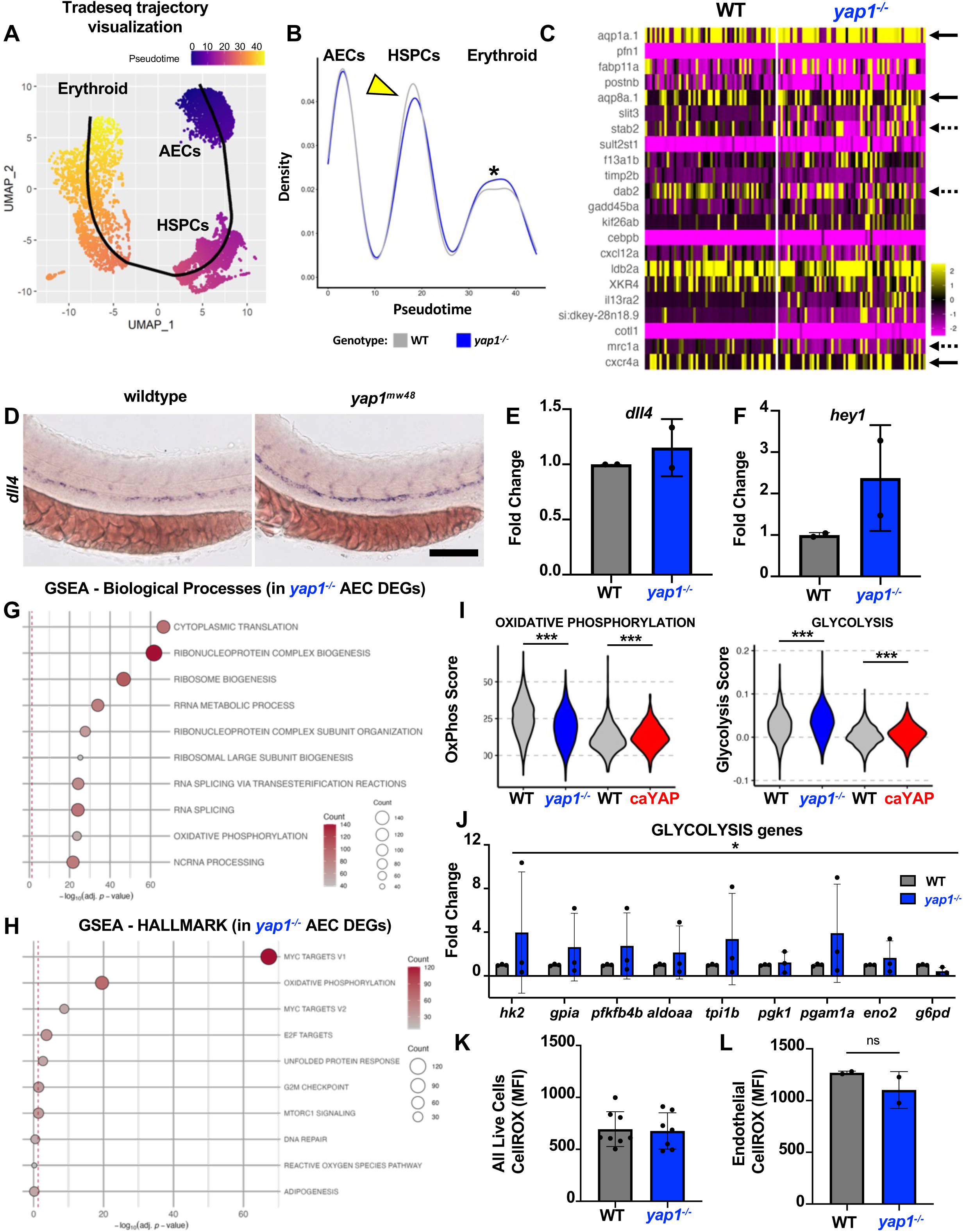
YAP loss causes EHT arrest by reversion to endothelial fates and blocked metabolic rewiring in HE cells. **A**) Pseudotime trajectory from Tradeseq algorithm across the EHT UMAP from arterial to erythroid identity, passing through HSPC intermediates. **B)** Proportion plot showing distribution of cells by genotype across the pseudotime trajectory from WT (grey) and *yap1^−/−^* (blue) cells from the LOF experiment. Note increased WT representation in HSPC peak compared to mutant (yellow arrowhead) and increased abundance of mutant cells in erythroid peak (*). **C)** Heatmap of top upregulated genes in *yap1^−/−^* HE cells compared to WT. Solid and dashed arrows denote arterial and venous associated genes, respectively. **D**) Whole-mount in situ hybridization (WISH) for *dll4* in trunk vasculature of WT and *yap1^−/−^* zebrafish at 30hpf (n ≥ 25 embryos per condition). Scale bar: 100um. **E, F)** Whole-embryo qPCR for *dll4* and Notch target gene *hey1* at 30hpf in WT and *yap1^−/−^*. Expression normalized to *18S* housekeeping gene (n = 2; Error bars indicate SD). **G)** Top GO:BP terms for Biological Processes associated with *yap1^−/−^* DEGs compared to WT in the arterial cluster. **H)** Top GO:BP terms for Hallmark Pathways associated with *yap1^−/−^* DEGs compared to WT in the arterial cluster. **I)** Violin plots for composite scores of Hallmark pathways (Oxidative Phosphorylation (left) and Glycolysis (right)) in the arterial endothelial cells (AEC) from YAP GOF and LOF experiments (n = 1156 WT, 818 yap1^−/−^ cells for LOF, 883 WT, 704 caYAP cells for GOF; unpaired Student’s t-test, ***p ≤ 0.001). **J)** qPCR on RNA from dissected tails of WT and *yap1^−/−^* embryos at 30hpf for a battery of glycolysis-associated genes. Expression normalized to *18S* housekeeping gene (n = 3; full panel evaluated by two-way ANOVA, *p<0.05; Error bars indicate SD). **K)** Mean fluorescence intensity (MFI) of CellRox-Orange (marker for reactive oxygen species (ROS)) in total live cells from 52hpf WT and *yap1^−/−^* embryos by FACS (n = 2; Error bars indicate SD). **L)** Mean fluorescence intensity (MFI) of CellRox-Orange (ROS) in *Tg(kdrl:EGFP)^s843^*endothelial cells from 52hpf WT and *yap1^−/−^* embryos by FACs (n = 2, unpaired Student’s t-test, n.s. = not significant. Error bars indicate SD).

Further inspection of the top upregulated genes in *yap1^−/−^ gata2b+* HE cells indicated a prominent number of arterial and venous markers compared to WT (**Figure 5C**). Visualizing a subset of these genes across the trajectory revealed noticeably higher levels of venous and arterial gene expression in the AECs of *yap1^−/−^*mutants (**Figure S5C**), suggesting a potential reversion to a pan-endothelial phenotype in the absence of Yap. In support of this interpretation, we observed increased *dll4* expression in the DA of *yap1^−/−^* embryos at 30hpf by WISH, similar to the published phenotype of *runx1^−/−^* embryos^87^ (**Figure 5D**). We corroborated these findings by whole-embryo qPCR for *dll4* and the Notch target gene *hey1* (**Figure 5E, F**). We also observed in the trajectory analysis that several Hox genes (including *hoxa9a*, a member of the HOXA cluster known to promote definitive hematopoiesis^88,89^) lagged behind WT levels in *yap1^−/−^* cells (**Figure S5D**), whereas the glutamine synthetase gene *glula,* a metabolic enzyme induced during erythropoiesis^90^, was elevated in *yap1^−/−^*AECs in the endothelial portion of the trajectory (**Figure S5E**) compared to WT. To get an unbiased perspective on key biological processes affected by Yap in arterial and HE cells, we performed GO enrichment on all differentially expressed genes (*yap1^−/−^* vs WT) across the entire AEC cluster. We found terms related to protein translation, cell growth/cycling, and oxidative phosphorylation (oxphos) represented in Biological Pathways and Hallmark annotations (**Figure 5G, H**). We then assessed the strength of the gene module score against Yap dosage for the following processes: mTorc1 signaling, heme metabolism, glycolysis, and oxphos. Oxphos and mTor signatures responded positively to Yap strength in a dose-dependent manner (**Figure 5I** and **S5F**), consistent with recent reports on hemodynamically-driven activation during EHT in mouse models^91,92^. Interestingly, glycolysis and heme metabolism were both elevated in *yap1^−/−^*AECs (**Figure 5I** and **S5G**), which may be reflective of an inability to shift from primitive to definitive blood forming potential; glycolysis genes, in particular, were measurably elevated in dissected tails from *yap1^−/−^* embryos by qPCR (**Figure 5J**). While there was no difference in the amount of reactive oxygen species (ROS) detectable in the bulk pool of live cells from whole dissagregated WT and *yap1^−/−^*embryos via FACS-based CellRox staining (**Figure 5K**), it was notably lower in the Kdrl+ endothelial population of *yap1^−/−^* mutants, suggestive of impaired oxphos in the endothelium, inclusive of HEC and newly specified HSPCs (**Figure 5L**). Together, these findings indicate that Yap is necessary for the correct metabolic rewiring of HE cells as they undergo EHT, and that reversion to endothelial identity and/or premature erythroid differentiation is a likely culprit for the reduced HSPC numbers observed in *yap1^−/−^* mutants.

### Comparison of YAP GOF/LOF and TAZ GOF transcriptomes identifies YAP-regulated metabolic hubs and TAZ-driven endothelial signatures in Runx1+ HE

Given our observation that TAZ possessed the ability to interact with RUNX1 and functionally compensate for YAP loss, we performed additional scRNA-squencing on sorted Kdrl+ WT and caTAZ endothelial cells to identify shared and divergent roles of the YAP/TAZ transcription factors in hematopoiesis. As with our YAP GOF and LOF experiments, we subsetted on the endothelial, hematopoietic, and erythroid clusters (**Figure S6A**). Our previous functional analyses indicated that YAP and TAZ overexpression can increase specification of *runx1^+^* cells in the DA, but only YAP provides the necessary transcriptional regulation to increase CD41^lo^ HSPC number. We therefore focused our bioinformatic analysis on *runx1+* HE, as this population represents a likely inflection point in HSC formation where divergent transcriptional programs could drive these differing phenotypes (**Figure S6B**). To compare the two overexpression models, we integrated the caYAP and caTAZ datasets **(Figure S7A, B)**, subsetted endothelial, HSPC, and erythroid populations, and confirmed cluster identity via EHT markers *gata2b, runx1,* and *cmyb* **(Figure S7C-E)**. We selected *runx1*+ HECs in the arterial cluster using MAGIC imputation and conducted GSEA analysis to identify cellular functions associated with gene expression changes caused by YAP and TAZ **(Figure 6A).** Broad trends included upregulation of migratory, glycolytic and nucleotide biosynthesis terms in the setting of caYAP (**Figure 6B-C**). In contrast, downregulated GO term themes in *runx1+* HE cells overexpressing YAP were largely associated with adhesion programs (**Figure S7F**), likely indicative of active remodeling, budding, and shifts in cell identity that characterize EHT. Notable upregulated terms in caTAZ cells included those associated with arterial formation and remodeling (**Figure 6D**), as well as developmental priming-related terms pertaining to WNT, retinoic acid (RA), FGF, and TGFβ signaling **(Figure 6E).** By contrast, downregulated terms in caTAZ *runx1+* HE cells were characterized by themes associated with hematopoietic migration as well as immune activation and inflammation (**Figure S7G**), previously shown to be required for HSPC production^93–97^. To more effectively compare our genesets to highlight drivers of our phenotypic results on HSPC formation, we generated a list of shared (regulated in the same direction, **Figure 6F**) and unique (regulated in the opposite direction, **Figure 6G**) GO terms from caYAP and caTAZ *runx1+* HEC, *gata2b+* HEC, and *runx1+* hematopoietic progenitors (see **Supplementary Table 1)**. Broadly, we delineated potential shared functions between YAP and TAZ in developmental priming, actin/cytoskeletal remodeling, and engagement of proliferative metabolic switches. Differentially regulated terms between the caYAP and caTAZ datasets generally correlated with activation of nucleotide biosynthesis **(Figure S7H)**, cell migration **(Figure S7I),** and downregulation of junction proteins **(Figure S7J**), suggesting that these hemodynamically responsive processes may be particularly influenced by YAP-specific transcriptional regulation to drive completion of EHT and HSPC production.

**Figure 6:**
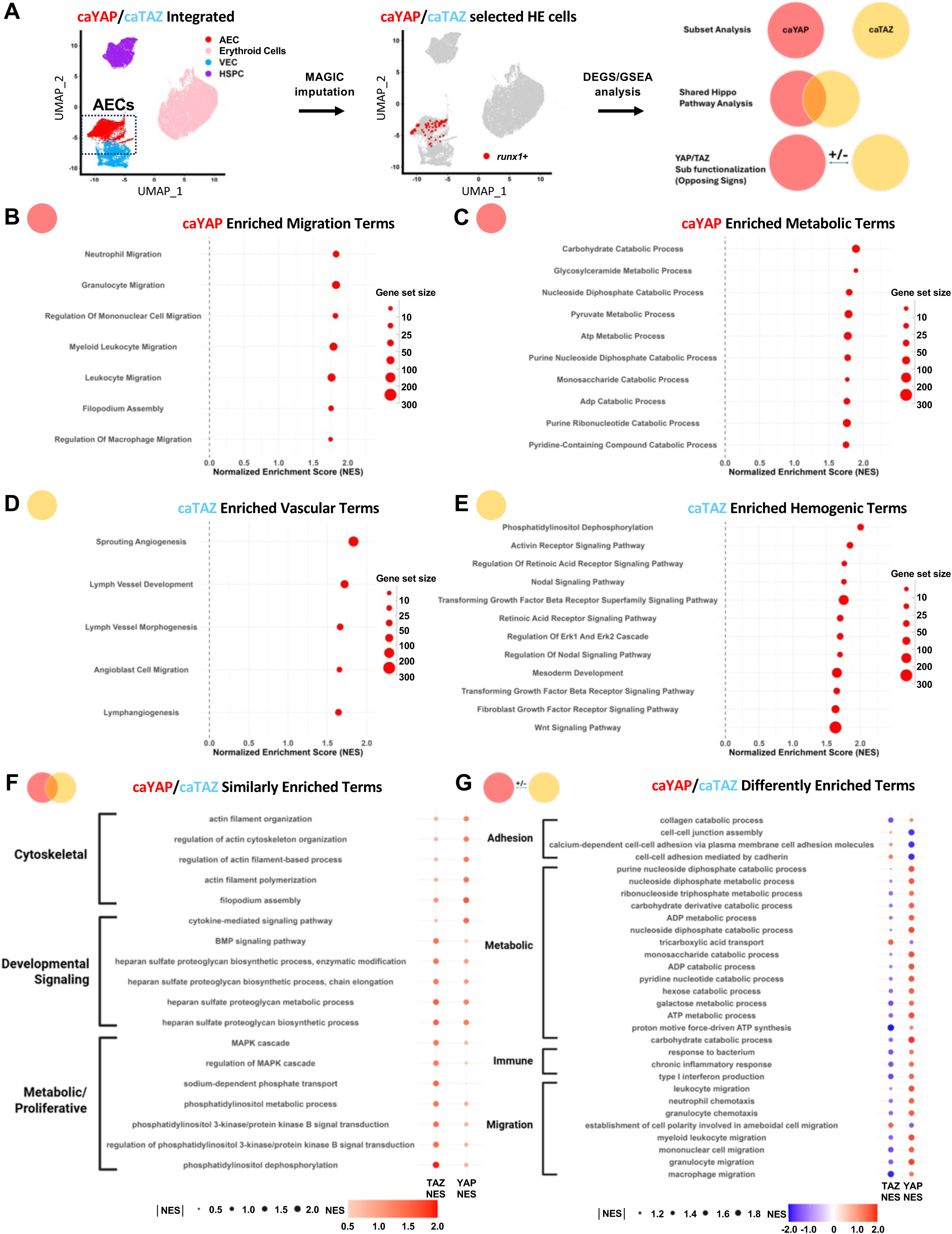
Sequencing of TAZ GOF EHT cells shows selective effect of TAZ on augmenting endothelial signature. **A**) Schematic denoting analysis pipeline and expression comparisons of caYAP vs caTAZ: (left) UMAP of AEC, VEC, Erythroid, and HSPC from the integrated caYAP/caTAZ dataset. (middle) MAGIC imputed *runx1*+ selected cells for downstream GSEA analysis. (right) GO:BP terms generated individually analyzed caYAP or caTAZ datasets (top), terms regulated in same direction (middle), and terms regulated in the opposite direction (bottom) between datasets. **B)** GO:BP terms reflecting enriched migration programs in the caYAP dataset. **C)** GO:BP terms reflecting enriched metabolic programs in the caYAP dataset. **D)** GO:BP terms reflecting enriched vascular programs in the caTAZ dataset. **E)** GO:BP terms reflecting enriched hemogenic programs in the caTAZ dataset. **F)** GO:BP terms reflecting cytoskeletal, developmental signaling, and metabolic proliferative programs upregulated in both caYAP and caTAZ datasets. **G)** GO:BP terms reflecting adhesion, metabolic, immune, and migration programs differentially regulated in the caYAP and caTAZ datasets.

### HSPC amplification by Piezo1 agonism is through YAP/TAZ convergence on proliferative signaling pathways

To define how these shared and divergent transcriptional programs influence HSC specification and guide functional validation, we examined select gene module scores identified in our analysis across all three genotypes: caYAP, *yap1^−/−^*, and caTAZ (**Figure 7A**). Interestingly, both caTAZ and *yap1^−/−^* cells maintained a high signature of cell-cell adhesion compared to caYAP. Cells exhibited a lower cell cycle signature in the context of YAP LOF, and increased nucleoside metabolism by YAP GOF. These observations suggest that retention of endothelial identity may underlie impaired EHT in both caTAZ and *yap1^−/−^* settings, with metabolic growth and migration powering HSPC production and expansion in YAP GOF. We next interrogated the cellular basis for reduced EHT in our caTAZ model. Prior studies have shown that Runx1 can regulate cell adhesion programs in HE early in EHT progression^98^. We hypothesized that the Runx1/TAZ interaction could hyperactivate this axis, potentially rendering a non-permissive environment towards productive EHT^99^: in support of this conjecture, we noted that the module score for the Reactome term ‘Transcriptional Regulation by Runx1’ was highly elevated in *runx1+* HE by caTAZ (**Figure S8A**) and confirmed that a panel of junctional and endothelial identity terms was significantly upregulated in caTAZ embryos via qPCR (**Figure 7B**). Live confocal imaging of Gata2b+ HE cells in triple transgenic *Tg(gata2b:Gal4^sd2^; UAS:Lifeact-EGFP^mu2^*^99^*; kdrl:TagBFP^mu293^)* reporter animals at 48hpf revealed a caTAZ-associated impact in shape and distribution of HE cells (**Figure 7C**). While standard quantification showed no difference in mean number of total Gata2b+ HE cells between WT and caTAZ conditions (**Figure S8B**), caTAZ expression altered the per-embryo spread of the data compared to WT (**Figure S8C**). Notably, the proportion of Gata2b+ HE cells that exhibited a flattened endothelial morphology, instead of the rounded budding phenotype characteristic of EHT, was nearly tripled in the cohort of caTAZ embryos examined (**Figure 7D**). These data collectively suggest that the failure to robustly expand HSPCs via caTAZ overexpression may be due to stronger activation/retention of endothelial programs that outweigh a positive impact on a hematopoietic signature, ultimately resulting in an inability to complete EHT.

**Figure 7:**
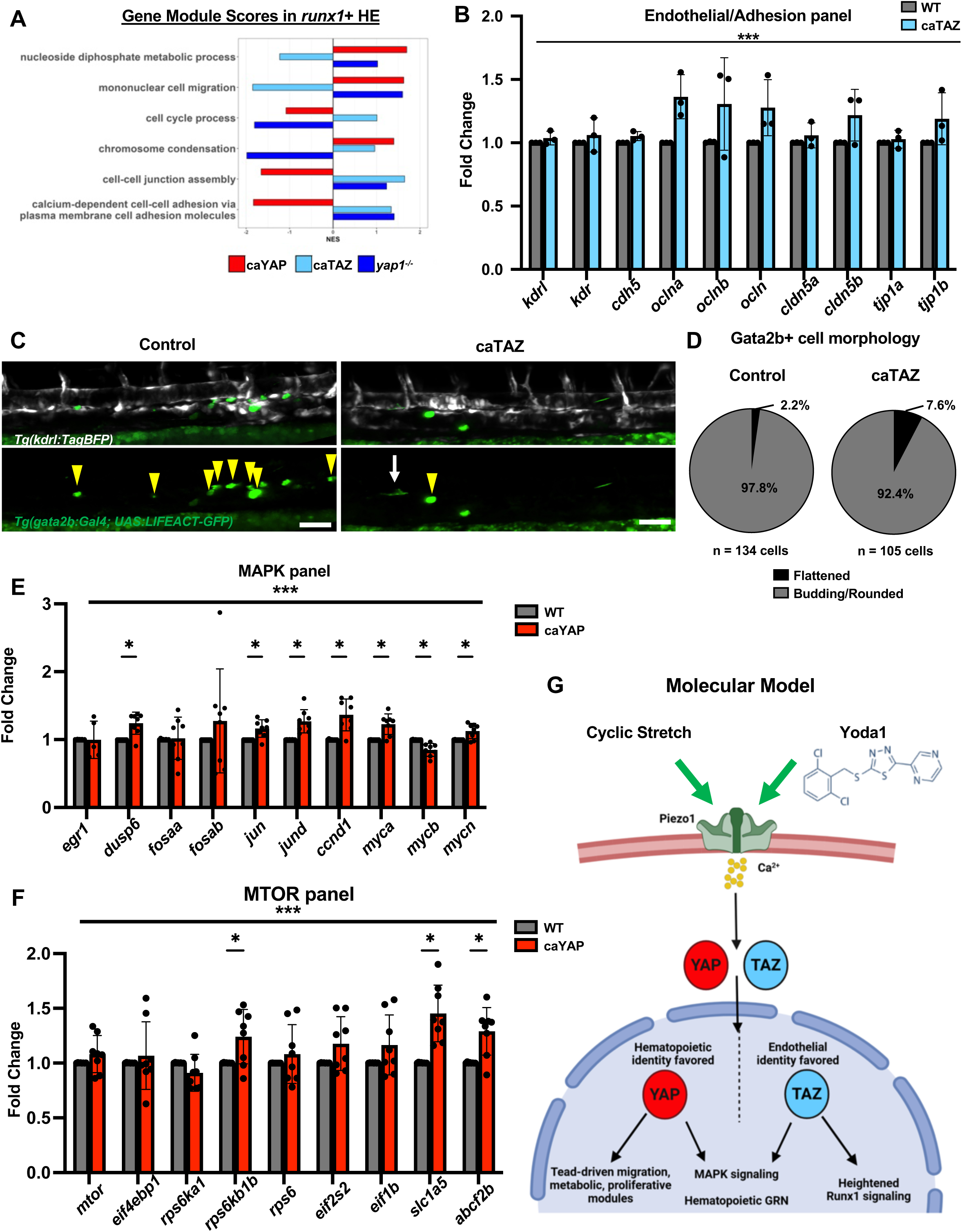
HSPC amplification by Piezo1 agonism acts via modulation of shared YAP/TAZ proliferation pathways. **A**) Gene module scores of select EHT processes in *runx1+* HE cells of different Hippo signaling genotypes (caYAP, *yap1^−/−^* and caTAZ) compared to WT. **B)** Whole-embryo qPCR for endothelial receptors and junction proteins at 30hpf in WT and caTAZ embryos following HS at 24hpf. Expression normalized to *eef1a* housekeeping gene (n = 3; full panel evaluated by two-way ANOVA, ***p<0.001; Error bars indicate SD). **C)** Maximum intensity projection of confocal z-stack of Gata2b+ HE cells (yellow arrowheads) in the DA at 48hpf in WT and caTAZ embryos. White arrow indicates HE cell with flattened, cobblestone morphology. Endothelial cytoplasm, HE cells visualized by *Tg(kdrl:TagBFP^mu293^)* and *Tg(gata2b:Gal4^sd32^; UAS:LIFEACT-GFP^mu271^)*, respectively. Scale bar: 50um. **D)** Quantification of Gata2b+ cell morphology in the VDA region from images in **C** (n = 134 cells from 25 WT, 105 cells from 23 caTAZ embryos). **E)** Whole-embryo qPCR for MAPK signaling pathway genes at 30hpf in WT and caYAP embryos following HS at 24hpf. Expression normalized to *18S* housekeeping gene (n = 8; full panel evaluated by two-way ANOVA, single genes by unpaired Student’s t-test *p<0.05, ***p<0.001 Error bars indicate SD). **F)** Whole-embryo qPCR for MTOR signaling pathway genes at 30hpf in WT and caYAP embryos following HS at 24hpf. Expression normalized to *18S* housekeeping gene (n = 8; full panel evaluated by two-way ANOVA, single genes by unpaired Student’s t-test *p<0.05, ***p<0.001 Error bars indicate SD). **G)** Molecular model of hemodynamic/pharmacologic activation of Piezo1 leading to YAP/TAZ activity in HE that controls shared and unique modules for cellular growth, hematopoietic lineage commitment and endothelial functions. Differential interactions with TEAD and the lineage-specific TF Runx1 allow mechanostimulation of YAP/TAZ to fine-tune EHT progression.

Because our functional and informatic analysis indicated that cell cycling and metabolic rewiring were key differences between caYAP and YAP LOF *runx1+* HE cells, and that a MAPK signature was enhanced by both YAP and TAZ in this population (**Figure 6F**), we explored the hypothesis that enhancing YAP activity either genetically (by caYAP overexpression) or pharmacologically (by stimulation of Piezo1) could increase HSPC amplification via metabolic-responsive proliferative signaling pathways. caYAP induction increased the expression of several MAPK family genes in zebrafish embryos, including cell cycle regulator *ccnd1* and transcription factors *myca*, *mycn*, *jun* and *jund* (**Figure 7E**). Likewise, cell cycle regulators, *ccnd1* and *cdc20,* were increased at 30hpf in zebrafish embryos exposed to Yoda1 (**Figure S8D**). Yoda1 treatment also led to more EDU+ cells in the DA at 48hpf (**Figure S8E**). YAP overexpression additionally drove upregulation of genes associated with mTOR signaling, including nutrient transporters (*slc1a5*, *abcf2b*) and the ribosomal protein S6 kinase *rps6kb1b* (**Figure 7F**). Blood flow-driven activation of these pro-growth pathways in HE by YAP could support the demands of proliferative expansion or migratory egress during EHT leading to increased HSPC production.

Taken together, our comprehensive transcriptomic and functional analysis of the role of YAP/TAZ transcription factors in HSPC development leads to the following cellular (**Figure S8F**) and molecular (**Figure 7G**) models. At the cellular level, YAP/TAZ function is required to maintain the specification and morphogenesis of a highly specialized endothelium-derived cell type, Runx1+ HE in the DA. TAZ overexpression increases hematopoietic GRN TF expression, but simultaneously amplifies an endothelial/adhesional signature in the same cells, physically blunting EHT progression. This may be due, at least in part, by over-stimulation or differential regulation of Runx1 transcriptional outputs mediated by TAZ/Runx1 synergy. By contrast, YAP overexpression preferentially drives TEAD-dependent metabolic and proliferative modules, in addition to regulation of the core hematopoietic GRN, while downregulating adhesion signatures, yielding increased EHT and HSPC production. Molecularly, Piezo1 is a mechanosensitive ion channel that can be biophysically (blood flow) or pharmacologically stimulated to activate YAP/TAZ signaling in the DA. Further dissection of the dual nature of YAP/TAZ regulation of endothelial and hemogenic gene programs may suggest strategies by which small molecule stimulation can be leveraged to preferentially promote efficient HSPC production *in vitro*.

## Discussion

While the requirement of hemodynamic cues for the production of HSPCs during embryonic development has now been well recognized, the mechanisms by which the sensing and transduction of those cues within HE fosters acquisition and/or commitment to hematopoietic fate has remained relatively undefined. Here, we show by transcriptomic and functional approaches in the zebrafish model that Yap and Taz transcriptional coactivators have an essential role in ensuring the full commitment of HE cells to EHT and HSPC fate by reifying a self-renewal hematopoietic GRN, amplifying Runx1 signaling, and boosting the metabolic and cycling capacity of the HE. In the absence of Yap, endothelial and erythroid-signature genes are upregulated in HE cells, suggesting these cells may experience a reversion to an endothelial fate or adopt primitive erythroid/EMP characteristics. This is reminiscent of a recent study highlighting Yap’s role in specifying distinct cell types within a tissue during developmental organ growth (cholangiocytes in the liver, and peripodial epithelium in the Drosophila eye)^100^. Regulation of general ‘pro-growth’ pathways as well as lineage-specific TFs help explain the Yap loss– and gain-of-function phenotypes observed in the zebrafish system.

Our finding that hemodynamic or pharmacologic stimulation of Piezo1 can activate the YAP/TAZ axis in human and zebrafish HE represents a significant step forward in providing a ‘membrane-to-nucleus’ mechanism for flow-directed initiation of EHT. Previous studies detailing the requirement for hemodynamics uncovered key secondary messengers downstream of WSS that promote HSC formation, including nitric oxide^34,36^, prostaglandin/cyclic AMP^101^ and Ca^2+^. Single-cell transcriptomics using mouse models in the absence of blood flow (such as the Ncx^−/−^ line) described impaired hematopoietic differentiation and oxphos deficiencies in HE cells^35^, without clearly delineated nuclear mediators for these effects. While our prior work established Rho-GTP/YAP mechanotransduction as an integrator of hemodynamics with EHT^40^, we did not identify membrane proteins that could relay biophysical stimuli. Cilia were previously described as a sensory organelle important for HSPC production in zebrafish^102^. Chemical activation of Piezo1 has been shown to promote EHT-derived HSPC production^55^, with more recent reports documenting that activation of Piezo1 influences proliferation and differentiation of mature murine HSCs, *in vivo*^103^ and *ex vivo*. Importantly, these effects appear to depend on the kinetics of Piezo1 activity: sustained chemical agonist treatment leads to differentiation and exhaustion, while transient induction promotes self-renewal and expansion. The hemodynamic profile in early embryonic development more closely resembles a pulsatile, heterogenous mechanical cue^104,105^, which may instruct downstream Piezo1/YAP output. The observation that the impact of the *piezo1^−/−^* phenotype in zebrafish is masked, at least in part, by the robust circulatory flow these embryos maintain highlights key features of the cell biological/environmental framework in which EHT occurs. WSS and CS never occur in isolation in the vertebrate AGM during the window of EHT; further work is needed to determine whether these forces interact redundantly or uniquely in HE. Likewise, while we have identified a role for Piezo1 in Yap activation, there are likely additional mediators of biomechanical stimulation that could act in a compensatory and/or collaborative manner. For example, integrins have been previously reported as mediators between extracellular forces and YAP mechanotransduction^75^; we and others have previously reported hematopoietic phenotypes for *itgb1b* loss-of-function in zebrafish^106,107^, but the extent to which this is connected to Yap signaling in HE and/or HSPCs remains unknown. Likewise, there may be as of yet undescribed contributions from other membrane ion channels and mechanosensors in HE that could serve to further fine-tune the precise level of YAP/TAZ regulatory activation across the DA.

The observation that Yap overexpression preferentially regulates a suite of metabolic pathways provides a previously undefined nuclear regulator able to couple a hemodynamic cue to the myriad changes needed in the switch from arterial endothelial to hematopoietic progenitor functionality. During sprouting angiogenesis in the retina, YAP/TAZ have been shown to enhance mTOR signaling to maintain cellular growth needed in proliferating vascular beds^108^. Our bioinformatics data shows a YAP-responsive mTOR signature, potentially in line with recent data linking mTOR to hemodynamically-driven EHT^92^. Curiously, one of our strongest YAP-driven metabolic signatures was ‘polyamine biosynthetic processes’; this has been documented in mouse liver and human cancer cells as a driver of growth^109^. YAP has further known roles in nucleotide biosynthesis^110^ and cell cycling^111^, which are critical in the formation of blood progenitors. A molecular axis connecting Piezo1/YAP to MAPK signaling has been described in proliferation in HeLa^59^ and other cancer cells^112^, and may be similarly controlling expansive growth during embryogenesis to establish the stem cell pool. Lastly, our finding that Yap appears to promote a glycolysis-to-oxphos shift in HE cells supports that this is indeed an actively regulated component of EHT. *Yap1^−/−^* endothelial cells have increased glycolytic gene expression despite relatively normal erythrocyte flow in the DA, strongly suggesting this is not an artefactual effect from hypoxia, but rather interrupted morphogenesis, associated with the failure to produce HSPCs.

The most unexpected aspect of our work is the differential functional outcomes from YAP and TAZ overexpression, and the implication of a tight regulation of signaling via the two TFs. While compensation/redundancy is a feature of YAP/TAZ in many tissue contexts, a growing body of work has identified unique functions of each^113^. Our findings that YAP drives EHT and HSPC production, while TAZ amplified an endothelial/migration signature is similar to the regulatory partition described for YAP and TAZ in lung cancer cells^114^. The basis for these effects will be of great interest for future work. One possible explanation is in the alternative DNA binding partners and complex formations that YAP and TAZ form within the tissue-specific milieu of TFs. For instance, crystal structures of TAZ/TEAD reveal a possible tetrameric configuration for TAZ and TEAD (in a 2:2 ratio) not observed with YAP^115^. Our complete GOF and LOF studies, together with functional studies of YAP/TAZ variants suggests that TEAD-directed activities play a primary role in the final numbers of HSPCs generated via EHT, and that dosage of YAP/TAZ is finely tuned during this process. In breast cancer, the interaction of RUNX1/3 with YAP limits the abilility of the latter to drive epithelial-to-mesenchymal transition^116^ (a process very similar to EHT), and we see observe RUNX1/TAZ synergistic activation of RUNX enhancers as seen in other contexts^49,50^. The ability of YAP and TAZ to interact with, and potentially alter/augment the transcriptional output of the master lineage regulator Runx1, therefore adds a layer of complexity to mechanically-regulated gene expression during EHT that must be systematically probed by functional analysis (and cannot be simply inferred by DNA binding motif presence). Indeed, the well-known Runx1+23 enhancer, a key element in the hematopoietic GRN, contains both TEAD and Runx1 motifs^117^, positioning it to be regulated by multiple YAP and/or TAZ DNA-binding complexes. Moreover, our findings that YAP and TAZ overactivation have, at times, opposing effects on hematopoietic and endothelial gene expression has significant implications for applications designed to stimulate these TFs for *in vitro* hematopoietic differentiation. Given that pharmacologic activation of Piezo1 would likely stimulate nuclear translocation of both TFs, it is highly probable that consideration of timing, dose, levels of YAP/TAZ and of TEAD/Runx1 DNA binding partners will all be necessary to balance the pro-hematopoietic and pro-endothelial arms of this mechanotransduction pathway to most efficiently stimulate not only enhanced specification, but completion of EHT and robust HSPC production for translational use.

## Acknowledgments

The authors would like to thank the Boston Children’s Hospital Flow Cytometry Core, HMS Single Cell Core, Harvard Biopolymers Facility, the BIDMC Confocal Core, the BIDMC and MCW Aquatics Facility staff and the Versiti Blood Research Institute Shared Resources Core Facility (RRID: SCR_025503) for their services, instrumentation, and specialist support. We are grateful to Chloe Baron for advice and training on scRNA-seq analysis, and thank Yi Zhou and Song Yang for raw data alignment and CellRanger processing. The *piezo1^um136^* line was generously shared by Nathan Lawson; we thank Brian Link, Didier Stainier and Leonard Zon for sharing many other zebrafish lines and reagents used in this work. We thank Alan Cantor, Stuart Orkin and Nan Liu for scientific input, and we are grateful to all members of the North, Daley and Schlaeger labs for scientific discussion and feedback on this study.

## Author Contributions

W.W.S and T.E.N. designed the project. W.W.S, S.G. and T.E.N. supervised the work and wrote the paper. W.W.S., S.G., Z.C.L., M.T.W., J.G., E. Molnar, L.R., Z.Y. and B.D.L. performed zebrafish experiments. S.G. and W.W.S. performed scRNA-seq data analysis. W.W.S. and Y.T. performed luciferase assays. R.M., C.L., A.M.T., O.S., K.K., R.J., C.K. and V.L. performed *in vitro* experiments with human CD34+ cells. W.Z. and C.A.M. generated and performed experiments with the piezo1-overexpressing zebrafish line. M.G.T. designed point mutant variants of TAZ and Runx1 for *in vitro* assays. E. Meader performed EDU assays, and M.T.W. performed CellRox assays. M.F. and M.A.N. provided code for bioinformatic analysis. G.Q.D. and T.M.S. provided reagents, protocols and support. All authors reviewed and edited the manuscript.

## Funding

This work was supported by: NIH K01DK129409 (WWS), TL1DK143273 (SG), R01 HL152636 (TEN) and RC2 DK120535 (GQD, TEN); and the American Society of Hematology (TEN).

## Disclosures

G.Q.D. holds equity and/ or receives consulting fees from Redona Therapeutics and iTCells, Inc.

## Methods

### Zebrafish Husbandry

Zebrafish were maintained in an IACUC-approved aquaculture facility at 28.5°C and utilized in accordance with approved animal protocols at Beth Israel Deaconess Medical Center, Boston Children’s Hospital, and the Medical College of Wisconsin. *yap1^mw48^*, *taz^mw49^* and *piezo1^um136^*alleles were identified by PCR amplification and restriction digest by TfiI, HinfI, and HpyCH4III, respectively. Band patterns are as follows: YAP genotyping [WT=305bp, MUT=157+141bp]; TAZ genotyping [WT=253+150+100+33bp, MUT=281+150+100bp]; Piezo1 genotyping [WT= 91+86+47bp, MUT= 128+91bp]. Primers and zebrafish lines are listed in the Key Resources table. Heat shock-inducible transgenes were activated at desired timepoints by incubation at 38°C for 45min.

### Chemical Exposures and WISH

Chemical treatments were performed in 6-well plates with ∼30 embryos/well. Controls were exposed to equimolar amounts of vehicle (DMSO). Piezo1 agonism was achieved with specified doses of Yoda1 (Cayman Chemical #448947-81-7). Heart rate was lowered with pimozide (Sigma # P1793). WISH experiments were performed according to published protocols with previously established probes^118^; embryos were fixed in 4% paraformaldehyde overnight at 4°C at the desired timepoint of analysis. *runx1* and *cmyb* WISH staining was quantitatively evaluated by categorizing staining intensity and coverage along the length of the dorsal aorta; typical patterns in control samples show a majority of embryos with a ‘beads-on-a-string’ pattern (scored as ‘medium’), with smaller numbers of individuals showing expanded clustering (‘high’) or patchy, low coverage (‘low’). All experimental conditions come from clutchmates obtained on the same day, and expression levels are binned with reference to the experimental control from that replicate. Scoring was performed by two independent operators to verify findings. Distribution plots show the total distribution of high/medium/low scoring embryos from all experiments (≥ 30 embryos per condition from at least 2-3 independent experiments, unless otherwise specified), and high-resolution images of a representative sample from the most prominent bin were captured on a Zeiss AxioImager microscope. Distributions were analyzed by Chi-square contingency tests to determine if deviations in experimental conditions were statistically significant from controls. The *piezo1* WISH probe was generated by antisense *in vitro* transcription from Horizon Discovery clone #EDR4422-213459652. Fluorescent WISH was performed with previously published protocols^119^. For live-imaging and fluorescent WISH experiments, embryos were exposed to 0.003% phenylthiourea (PTU) in embryo medium to prevent pigment formation. Confocal imaging was performed on a Zeiss LSM880 microscope, with live embryos anesthetized in 168ug/mL tricaine and immobilized in 1% low melting point agarose to avoid movement, as described^120^.

### Zebrafish Morpholino Injections

1-cell stage oocytes were injected with equal amounts standard control, 4ng *silent heart/tnnt2a*^121^, 5ng *yap1*^122^ or 1ng *taz*^123^ morpholinos obtained from Gene Tools (sequences in Key Resources table).

### Transgenic Lines

All zebrafish lines generated in this study were made via standard Gateway cloning and Tol2 transgenesis techniques, using plasmids containing full-length zebrafish ORFs for YAP1 (Uniprot: Q1L8J7) and TAZ (Uniprot: Q307I6). The QuikChange Mutagenesis II kit was used to make YAPS54A and TAZS48A point mutations on entry vectors for the YAPS87A and TAZS79A ORFs, respectively. The TAZS79A variant was introduced into an entry vector for WT TAZ. All variants in this study were recombined into the pDestTol2 destination vector with the following entry vectors: p5E-Hsp70l, pME-mCherry-NoStop, p3E-p2A-(YAP/TAZ variant). A single FLAG tag was added in-frame at the N-terminal portion of each variant. Single-cell oocytes were injected with ∼50pg DNA and ∼60pg Tol2 transposase RNA for genomic integration. Offspring from F0 adults were selected based on mCherry expression following heat shock to obtain the F1 generation, and all experiments were performed in F2 or F3 embryos.

### Confocal Microscopy

Live-imaging of zebrafish embryos was performed on Zeiss LSM880 and Nikon AXR confocal microscopes. Embryos were treated with 0.003% PTU in embryo medium to block pigment formation, and anesthetized with 168 ug/mL tricaine to prevent gross embryo movement. Embryos were immobilized in 1% low melting point agarose on glass bottom dishes for duration of imaging.

### EDU Assay

To visualize proliferating cells in the dorsal aorta, embryos were incubated for 1hr on ice in 500 mM EDU/ 10%DMSO in embryo medium starting at 30hpf, allowed to recover for 1hr at 28.5°C and fixed at 32hpf. The Molecular Probes Click-iT EdU Cell Proliferation Kit for Imaging (Catalog #: C10340) was used to perform detection and imaging of EDU incorporation.

### Flow Cytometry Analysis – Zebrafish Cells

For all FACS experiments, groups of 5-12 dechorionated embryos were collected in 1.5m L Epi tubes, rinsed with PBS and resuspended in 500uL PBS/1mM EDTA containing 75ug/mL Liberase (Sigma 5401119001). Dissociation to single cell suspension was achieved by incubation at 34°C with periodic mechanical disruption for at least 30min. Cell suspension was filtered through a 30um filter into 5mL FACS tube, and the liberase quenched with 2%FBS/PBS. Cells were centrifuged at 400g for 5min at 4°C, and the pellet resuspended in 2%FBS/PBS supplemented with 1:1000 dilution of Sytox Red live/dead stain (Invitrogen # S34859). Unlabeled and single-transgenic embryos served as gating controls. CD41:GFP^lo^ HSPCs were gated as previously described^124^. To measure oxidative stress, CellROX Orange (Invitrogen #C10443) was added to fresh live cell suspensions 30min before analysis at a 1:1000 dilution.

### Single Cell RNAseq Sample Prep

For scRNA-sequencing, endothelial cells were FAC-sorted from dissected embryo tails at 36hpf. Insulin needles were used to remove the tail from the head and majority of the yolk ball; ∼400 tails per condition were collected into a single 1.5mL tube. For the YAP/TAZ GOF experiments, *Tg(Hspl70:mCherry-p2A-Flag-YAP1^S87A^)* or *Tg(Hspl70:mCherry-p2A-Flag-WWTR1^S79A^)* adults were crossed to *Tg(kdrl:EGFP)^s843^* fish. GFP+ embryos were given a 38°C x45min heat shock at ∼14hpf, and the resulting mCherry – and mCherry+ clutchmate samples were processed for FACS and scRNA-seq library prep at 36hpf. For the YAP LOF experiments, large group crosses of AB (WT) and homozygous *yap1^mw48^−/−* adults carrying the *Tg(kdrl:mCherry)^s916^* transgene were performed to non-transgenic adults of the same genotype to obtain large numbers of stage-matched wildtype and *yap1^−/−^*embryos. Samples were processed for FACS and scRNA-seq library prep at 36hpf. FACS sample prep was performed exactly as described, with the exception that cells were sorted directly into 10%FBS/PBS. Samples were transferred to the Harvard Medical School Single Cell Sequencing Core for 10x Genomics library preparation. Target cell recovery was ∼10,000 cells per sample; sequencing was performed at the Harvard Biopolymers Facility on an Ilumina NovaSeq 6000 to an approximate depth of 30,000 reads/cell. Each GOF and LOF experiment was performed twice.

### scRNA-seq Bioinformatic Analysis

Briefly, raw data from sequencing was trimmed to remove adaptors and aligned to a custom version of the zebrafish genome corresponding to GRCz11 containing sequence of transgene fluorescent proteins used in the experiments. The 10x CellRanger pipeline was used to assign reads to cells. The SoupX algorithm^125^ was used to estimate and remove the contamination fraction of reads from all samples, using a Rho value between 2-5%. All bioinformatics procedures for quality control, filtering, clustering and UMAP visualization were performed in RStudio (ver 4.2.3) using Satija Lab Seurat (v4.3.0.1) functions. Cells were filtered using the following cutoffs: nUMI >500, nGene>600, mitoRatio <0.2. An integration step was used to merge the control/experimental samples from the YAP GOF and LOF experiments. Unbiased clustering was performed at a resolution of 0.4, and identities assigned using previously published datasets^77^. Endothelial and hematopoietic clusters of interest were subselected and reclustered using the same strategy to profile EHT populations. Similar cell types within large clusters were collapsed to reflect broad populations, and the SCT assay was used for all FeaturePlot() visualization. Gata2b levels from the integrated assay were used to establish an HE cutoff value of 3. The AddModuleScore() function from the Seurat package was used to measure the effect of YAP dosage on literature-based curations of hematopoietic TFs and YAP target genes, as well as the HALLMARK gene sets for glycolysis, oxidative phosphorylation, mTOR signaling and Heme Metabolism. Differential gene expression analysis was performed using the FindMarkers() function with default settings, comparing YAP GOF and LOF samples to their respective WT controls.

### scRNA-seq Trajectory Analysis

To interrogate differential gene expression over the course of a developmental trajectory dependent upon genotype covariates, we utilized the trajectory analysis packages Condiments^126^ and Monocle3^127–129^. Utilizing the standard Condiments/slingshot/tradeseq workflow, conditions were ascribed to the genotype of comparison: either caYAP vs WT or *yap1^−/−^* vs WT. Utilizing slingshot, each genotype was fitted to a pseudotime trajectory. Differential progression analysis was performed to assess differences in cell state across trajectories. The top 10000 variable genes across genotype were fit to a differential expression model using Tradeseq’s fitgam function, ranking differentially expressed genes for subsequent visualization. We compared these models to those created via the Monocle 3 workflow. Seurat objects were converted to CDS objects using the Satija lab’s Monocle 3 Seurat wrapper. Trajectories were fitted to UMAP space using both genotypes. Each genotype condition was then subset for visualization of gene expression. We selected genes we identified as differentially expressed either through DEG analysis or through the Condiments workflow for visualization in Monocle3. Gene expression analysis was performed in Monocle 3 over the course of a trajectory and each model’s expression values were superimposed on an expression plot in ggplot for ease of comparison.

### Zebrafish qRT-PCR analysis

RNA was extracted from pooled whole embryos or dissected tails (30 embryos/condition) using phenol/chloroform extraction. The aqueous layer was run through Qiagen Micro RNeasy columns to recover RNA, and 1ug of total RNA was used to generate cDNA using the BioRad iScript cDNA synthesis kit (BioRad). qRT-PCR was performed on a Bio-Rad CFX384 in 384-well plates using Sybr Green detection, with samples run in duplicate or triplicate. All experiments were done with ≥3 biological replicate pools/condition using the primers listed in the key resources table. Expression was normalized to a recommended primer pair for the zebrafish 18s housekeeping gene^130^, unless otherwise stated.

### Luciferase Assays

Luciferase assays were performed with expression vectors for full-length human transcription factor variants for YAP/TAZ obtained from Addgene (ID numbers in Key Resources table). A V5 tag was added to the C-terminus of a human RUNX1 expression plasmid in-house via standard cloning techniques. The QuickChange Mutagenesis II kit was used to introduce point mutations in this study not available on Addgene. 6xOSE-LUC was a gift from Dr. Stephen Weiss (U. of Michigan), and the SV40-RenillaLUC was a gift from Dr. Andrew Lassar (Harvard Medical School). All experiments were performed with the Promega Dual Reporter Luciferase system. HEK293T cells were seeded at ∼6-8K cell/well in 96 well plates, and allowed to attach overnight. Cells were then transfected with Lipofectamine and Plus Reagent with each well receiving: 250ng FireflyLUC, 10ng SV40-RenillaLUC, and 250ng of each TF DNA. pBluescript II KS(+) was used as a normalizer to keep DNA amount constant in the case of single TF conditions. After 24hrs, cells were lysed in 30uL Passive Lysis Buffer, and 20uL of lysate added to 100uL LARII reagent in a BioTek Plate Reader to record luminescence. The ratio of Firefly/Renilla luciferase activity was used to normalize for cell viability/transfection efficiency.

### iPSC Derivation and Maintenance

Human iPSCs were maintained on Matrigel matrix (Corning, 354277) using Stemflex media (Gibco, A3349401) in 5% CO^2^ at 37° C and passaged as clumps using ReleSR^TM^ (Stem Cell Technologies). Lines used include WT #1157.2 (generated from healthy donor PBMCs at Boston Children’s Hospital under IRB protocol #09-02-0068^29,131^).

### Human CD34+ Hemogenic Endothelial Cell Culture

120k cells were seeded onto a 10cm plate coated with matrigel via clump passaging. CD34+ hemogenic endothelium was generated using the STEMdiff Hematopoietic Kit (Stemcell technologies, 05310) according to manufacturer’s instructions until day 7, when hematopoietic cells begin to emerge. At this point, cells were dissociated using Accutase for 3-4 minutes at 37° C, quenched in flow buffer (2% FBS in PBS), and passed through a 40uM filter preceding flow cytometric analysis and CD34+ magnetic cell isolation (MACS). For MACS, cells were incubated for 30 min with magnetic bead coupled antibodies against CD34 in the presence of blocking reagent at 4°C with gentle rocking, followed by a wash step, then passed through a primed LS column. CD34+ enriched cells were seeded onto a Matrigel-coated 48 well at (50k/well) in EHT media as described previously^40^. EB media was supplemented with 10 ng/ml BMP4, 5 ng/ml FGF2, 15 ng/ml VEGF, 10 ng/ml IL6, 5 ng/ml IL11, 25 ng/ml IGF1, 50 ng/ml SCF and 2 units/ml EPO.

### Yoda1 Treatment of CD34+ Colonies

24 hours post plating, floating cells were washed away with PBS and fresh EHT media was added (500ul/48w). Then, cells were treated with 0.5uM Yoda1/DMSO. 24 hours post treatment, cells were harvested for RNA extraction via the RNeasy Plus Micro kit (Qiagen) for qPCR analysis.

### Modified Loh Protocol for HLF-reporter Detection and Hematopoietic Progenitor Quantification

Human pluripotent stem cells were cultured in mTeSR Plus media (Stem Cell Technologies #100-0276) with 1% Penicillin/Streptomycin (Corning # 30-002-CI) on tissue culture-treated plates coated with 1% Geltrex matrix solution (Gibco #A1413301) in DMEM/F-12 (Thermo Fisher #36254). Cells were clump passaged every 3-5 days using Versene (Thermo Fisher #15040066. Medium was replenished every 1-2 days. hiPSCs were differentiated into CD34+CD45+ hematopoietic progenitors within ten days as described previously^31^. Additionally, Zheng et al., 2025 provides detailed protocols for differentiation^132^. Briefly, hPSCs were dissociated using Accutase (Thermo Fisher # 00-4555-56), and seeded as single cells. iPSCs were induced to form posterior primitive streak within 48 hours, followed by formation of lateral mesoderm in 24 hours, and subsequently arterial endothelium in another 24 hours. Cells were subsequently dissociated into single cells using Accutase and re-plated at high density (∼500,000 cells/cm^2^) on plates coated with vitronectin (Thermo Fisher #A14700) and wild-type DLL4. Cells were then induced to form hemogenic endothelium for 72 hours, followed by induction of hematopoietic progenitors.

The Piezo1 ion channel agonist Yoda1 (Cayman #21904) was added at 0.5uM from Day 5-7 of differentiation, corresponding to specification of hemogenic endothelium from artery endothelium. Alternatively, DMSO was added as a vehicle control.

### Flow Cytometry Analysis – Human iPSC-derived Cells

Cells were collected on the final day of differentiation after 10-minutes of dissociation with Accutase at 37C. Accutase was diluted with FACS buffer (PBS + 2% FBS [Stem Cell Technologies #100-0180]), followed by centrifugation at 300G for 5 minutes at room temperature. Prior to staining, cells were passed through a 40µm. Fluorophore-conjugated antibodies were added to cells in FACS buffer, specifically: BV786 mouse anti-human CD34 (Clone: 581; BD Biosciences #743534) and FITC mouse anti-human CD45 (Clone: 2D1; BD Biosciences #347463). Cells were stained on ice for 30 minutes, shielded from light. After staining, cells were washed twice with FACS buffer and resuspended in FACS buffer with DAPI (1:100) for viability discrimination. Flow cytometry was performed on a Sony MA900 Multi-Application Cell Sorter. Data analysis was performed in FlowJo.

### Immunohistochemistry

Following Yoda1 treatment, cells were fixed with 4% PFA at room temperature for 10 mins. Cell permeabilization was carried out using 0.5% Triton-X-100 in PBS for 10 min at 4°C. Cells were then treated with blocking buffer (2% BSA in 0.1% Triton-X-100 in PBS) for 30 mins at 4°C. Primary antibody (1:50 dilution in blocking buffer, YAP SC-101199, Santa Cruz Biotechnology) staining was carried out for overnight at 4C. Subsequently, cells were washed with blocking buffer (3x, 10 mins each) and treated with secondary antibody (1:450 dilution in blocking buffer, Alexa Fluor-647 anti-mouse IgG, Jackson ImmunoResearch #715-605-150) and AlexaFluor-488 Phalloidin (1:100 dilution in blocking buffer, Invitrogen #A12379) for 1 h at 40°C. Finally, cells were treated with Hoechst (1x, 10 mins), and washed with blocking buffer twice before confocal imaging using a Leica SP8 machine.

### Quantification and Statistical Analysis

Unless otherwise specified, all experiments were performed in 2-3 independent biological replicates. Statistical analysis was performed with GraphPad Prism, and graphs were generated with Prism and ggplot2 (RStudio). Unpaired two-tailed Student’s t-test or one-way ANOVA were used to determine if differences in means between two or more groups, respectively, were statistically significant (*P* < 0.05 or lower); Chi-squared analysis was used to evaluate deviations in distributions between control and experimental groups of embryos from WISH analysis. Two-way ANOVA analysis was performed on qPCR gene panels to evaluate if experimental treatments were correlated with expression changes of the whole panel.

## Data Availability

The raw data from scRNA-sequencing experiments associated with this study has been uploaded to the Gene Expression Omnibus repository with the accession number GSE335024. All relevant code and supporting data are available from the authors upon reasonable request.

---

**Supplemental Figure 1:**
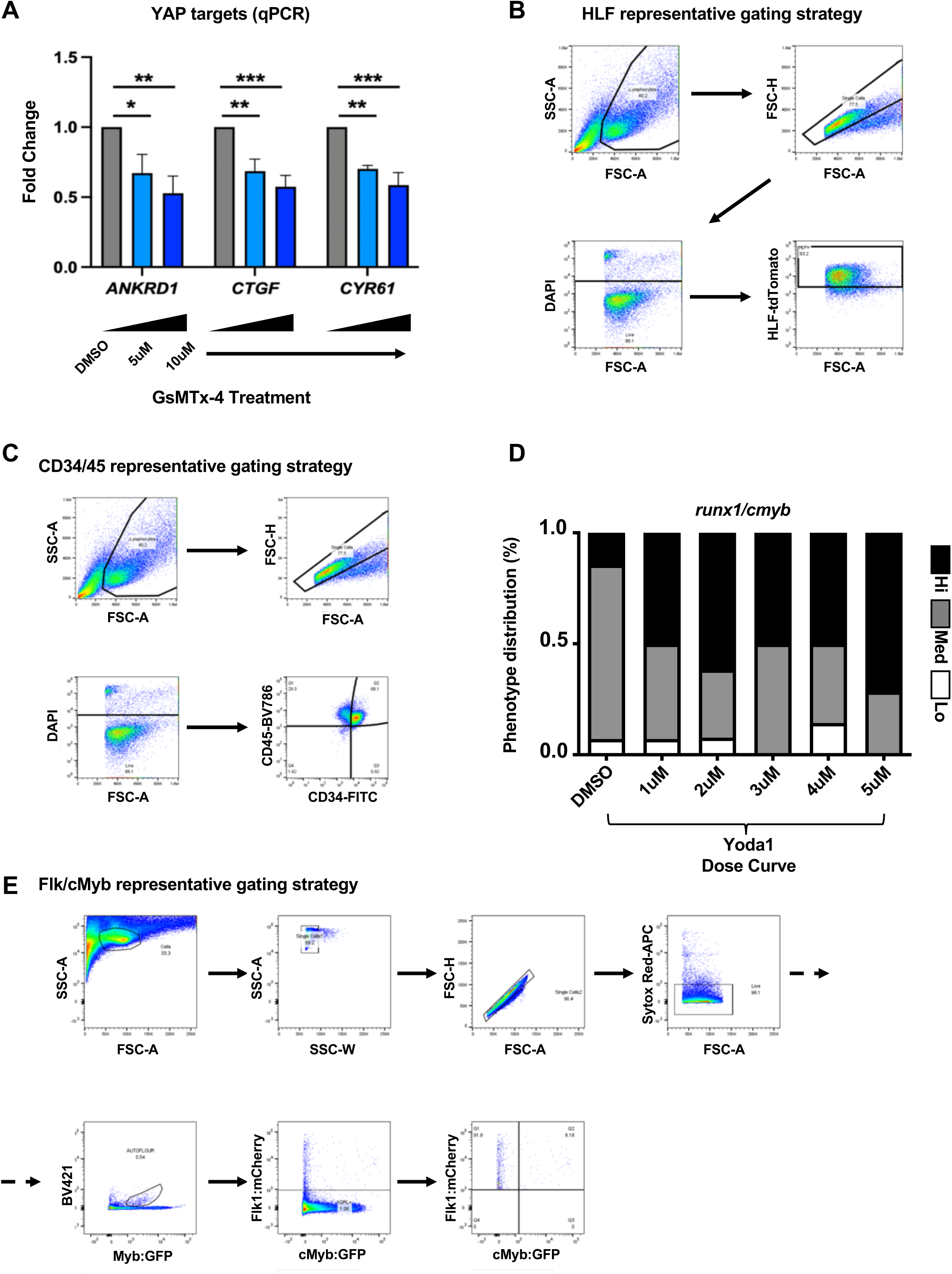
Piezo1-YAP axis can be pharmacologically stimulated to drive HSPC production in human HE cells and zebrafish. **A**) qPCR for human YAP target genes in human CD34+ cells exposed to DMSO or GsMTx-4 for 24hrs. Expression normalized to *GAPDH* housekeeping gene (n = 3, one-way ANOVA, *p ≤ 0.05, **p ≤ 0.01, ***p ≤ 0.001). **B)** Flow cytometry gating strategy for HLF-tdTomato reporter. **C)** Flow cytometry gating strategy for human CD34 and CD45 expression. **D)** Qualitative distribution plot for *runx1/cmyb* expression WT embryos exposed to DMSO or a dose curve of Piezo1 agonist Yoda1 from ∼10ss-36hpf (n ≥ 13 embryos/condition). **E)** Flow cytometry gating strategy for quantifying zebrafish Flk+/cMyb+ HSPCs in *Tg(kdrl:mCherry^s916^; cmyb:EGFP^zf169^)* transgenic embryos.

**Supplemental Figure 2:**
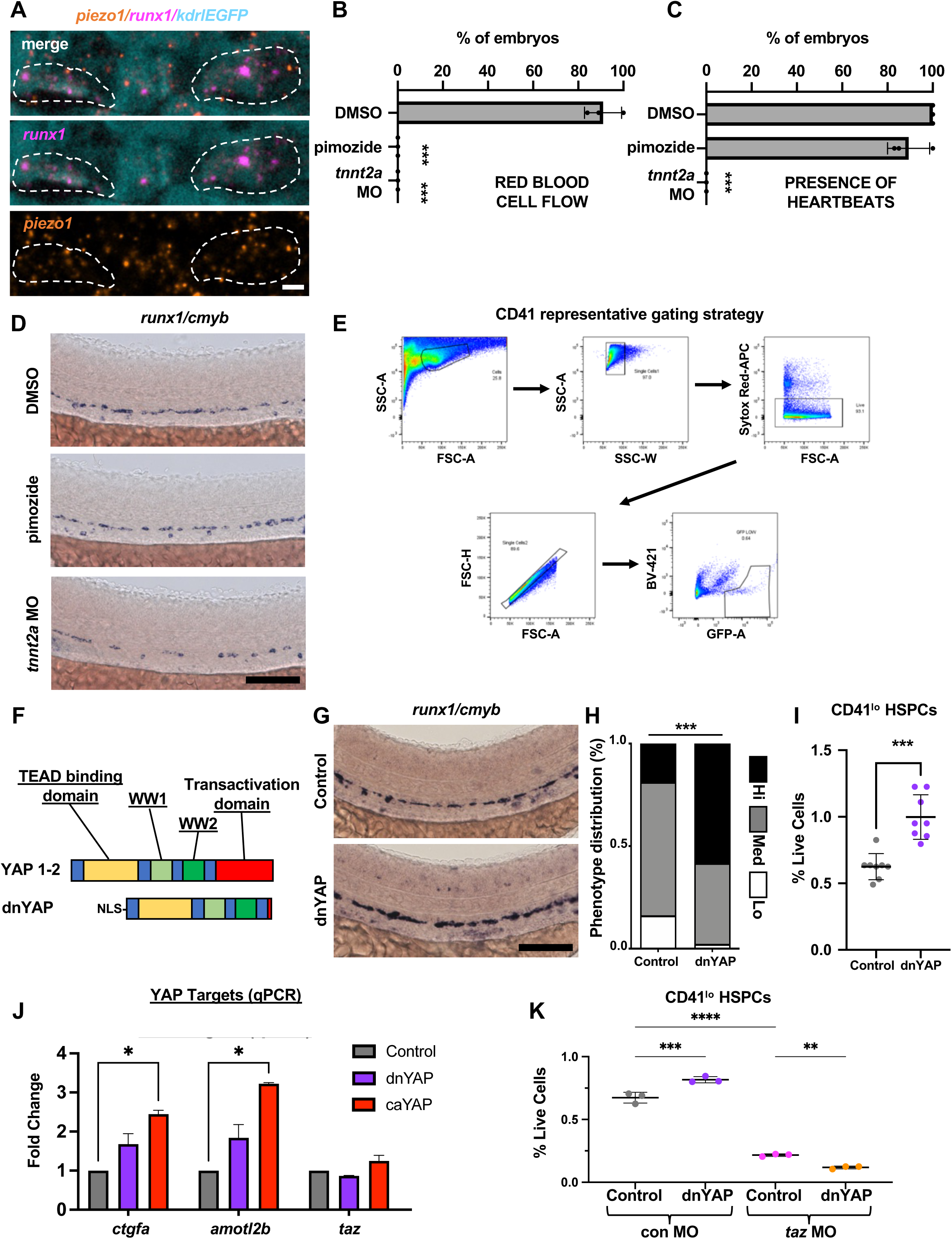
Piezo1 relays cue from hemodynamic stretch to YAP/TAZ signaling. **A**) Single slices of confocal z-stack of fluorescent in situ showing colocalization of *piezo1* mRNA (orange) in *runx1+* hemogenic endothelial cells (magenta) in the dorsal aorta (cyan) at ∼32 hpf. Vasculature is labeled by transgenic GFP expression in *Tg(kdrl:EGFP)^s843^* embryos captured with anti-GFP antibody staining. Scale bar: 5um. **B)** Quantification of red blood cell circulation (proxy for presence of wall shear stress) in the dorsal aorta at 36hpf. No RBCs/WSS occurs in both 4ng *tnnt2a* MO and 25uM pimozide treated embryos (n = 3, unpaired Student’s t-test ***p ≤ 0.001; Error bars indicate SD). **C)** Quantification of heartbeat presence (proxy for source of cyclic stretch-generating pulses) at 36hpf. 25uM pimozide embryos maintain irregular heart contractions, while 4ng *tnnt2a* MO embryos have none, resulting in a ‘stretch only’ force profile (n = 3; unpaired Student’s t-test ***p ≤ 0.001; Error bars indicate SD). **D)** WISH for *runx1/cmyb* shows relatively normal levels of hematopoietic cell production in 36hpf embryos exposed to 25uM pimozide (24-36hpf), compared to decreases in 4ng *tnnt2a* MO (n ≥ 50 embryos/condition). Scale bar: 100um. **E)** Flow cytometry gating strategy for quantifying zebrafish CD41^lo^ HSPCs in *Tg(itga2b:EGFP^la2^)* transgenic embryos. **F)** Schematic for design of heat shock-inducible dominant-negative YAP (dnYAP) construct. **G, H)** WISH for *runx1/cmyb* in control and dnYAP embryos at 30hpf following HS at 10ss. Qualitative phenotypic distribution plot of embryos scored with low, medium, or high *runx1/cmyb* expression in the VDA (n = 43 control, 43 dnYAP; chi-squared test ***p ≤ 0.001). Scale bar: 100um. **I)** FACs analysis of CD41:EGFP^lo^ HSPCs at 72hpf in control and dnYAP embryos (n = 10, unpaired Student’s t test ****p ≤ 0.0001; Error bars indicate SD). **J)** Whole-embryo qPCR for YAP target genes *ctgfa* and *amotl2b* and the YAP paralogue *taz* at 30hpf in control, dnYAP or caYAP embryos following HS at 24hpf. Expression normalized to *eef1a* housekeeping gene (n=2; Error bars indicate SD). **K)** FACs analysis of CD41:EGFP^lo^ HSPCs at 72hpf in control and dnYAP embryos injected with 8ng standard control or *taz* MO with a HS administered at 24hpf (n = 3, unpaired Student’s t test **p ≤ 0.01, ****p ≤ 0.0001; Error bars indicate SD).

**Supplemental Figure 3:**
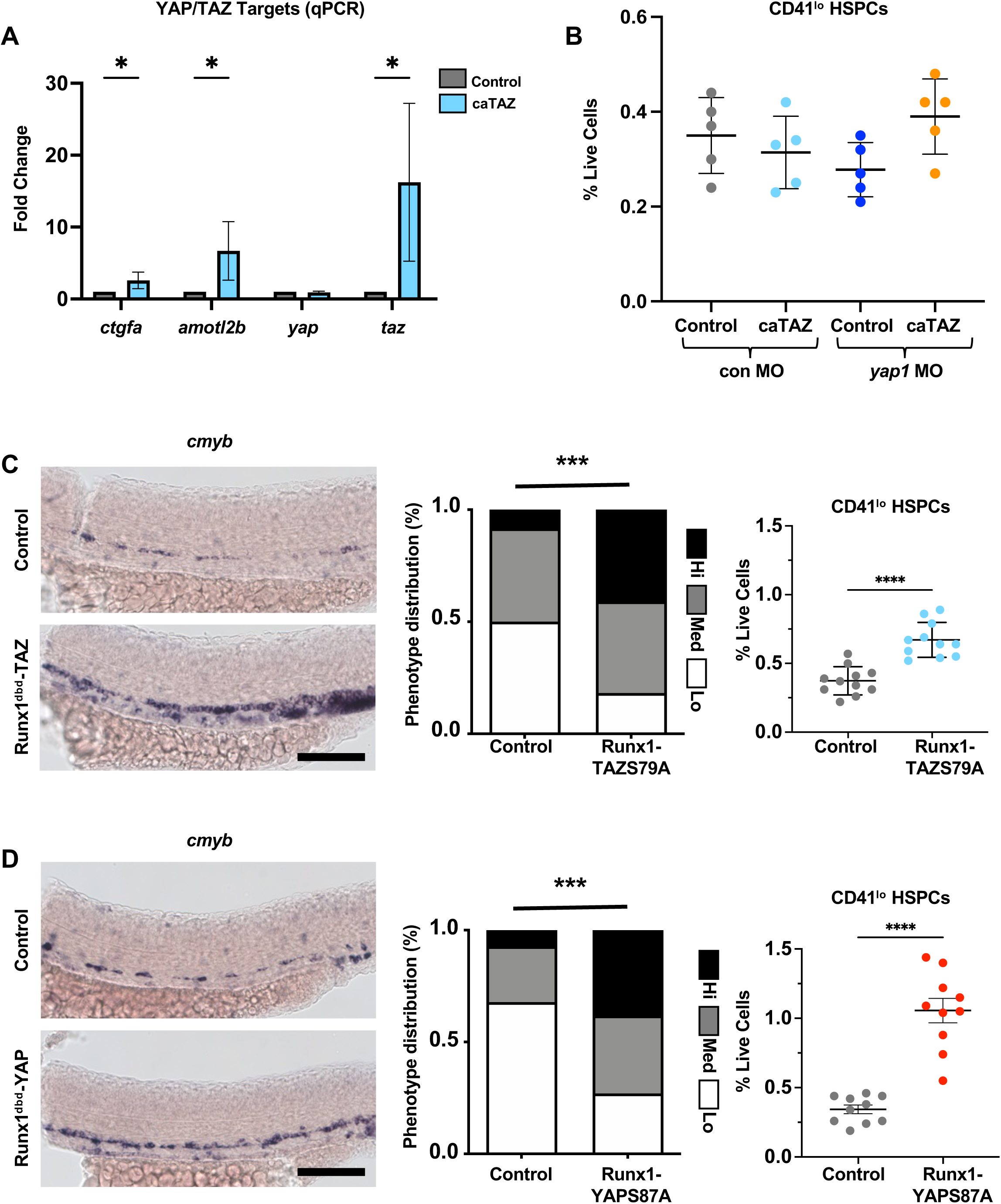
Temporal control of YAP deficiency demonstrates functional role for WWTR1/TAZ paralogue during EHT. **A**) Whole-embryo qPCR for YAP target genes *ctgfa* and *amotl2b*, and the YAP paralogue *taz* at 30hpf in control or caTAZ embryos following HS at 24hpf. Expression normalized to *eef1a* housekeeping gene (n = 8; Error bars indicate SD). **B)** FACs analysis of CD41:EGFP^lo^ HSPCs at 72hpf in control and caTAZ embryos injected with 5ng standard control or *yap1* MO with a HS administered at 24hpf (n = 2; Error bars indicate SD). **C)** WISH for *cmyb* expression with qualitative phenotypic distribution plot at 30hpf, and FACs analysis of CD41:EGFP^lo^ HSPCs at 72hpf in control and HS:Runx1^dbd^-TAZS79A embryos (synthetic tethering of TAZ to Runx1 enhancers). HS administered at 24hpf (n ≥ 20 embryos/condition for WISH, chi-squared test *p ≤ 0.05; n ≥ 11 for FACs, unpaired Student’s t-test ****p ≤ 0.0001; Error bars indicate SD). Scale bar: 100um. **D)** WISH for *cmyb* expression with qualitative phenotypic distribution plot at 30hpf, and FACs analysis of CD41:EGFP^lo^ HSPCs at 72hpf in control and HS:Runx1^dbd^-YAPS87A embryos (synthetic tethering of YAP to Runx1 enhancers). HS administered at 24hpf (n ≥ 50 embryos/condition for WISH, chi-squared test ***p ≤ 0.001; n ≥ 10 for FACs, unpaired Student’s t-test ***p ≤ 0.001; Error bars indicate SD). Scale bar: 100um.

**Supplemental Figure 4:**
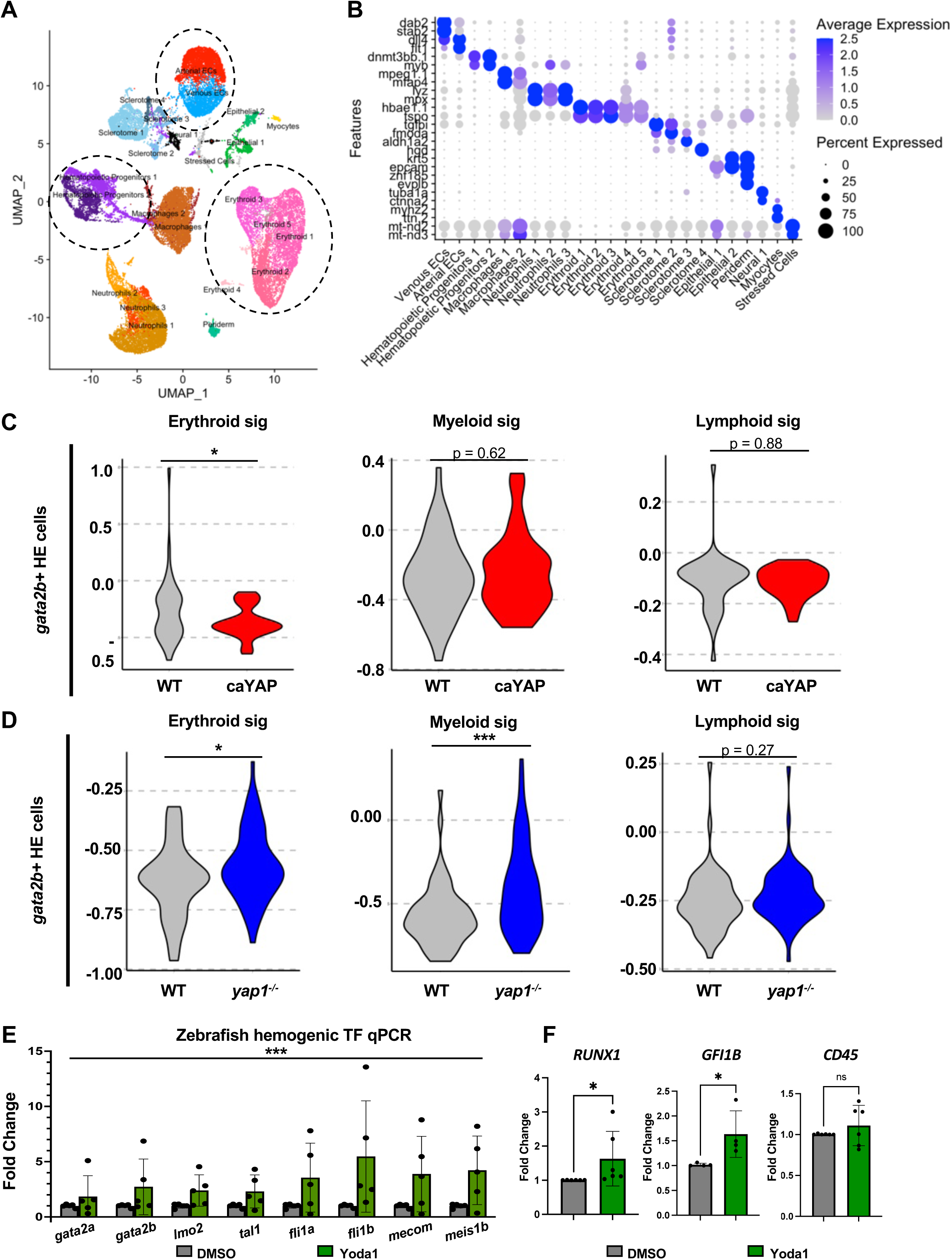
YAP promotes self-renewal and stemness in early hemogenic endothelium. **A**) UMAP of all combined integrated GOF and LOF Kdrl+ cells. Dashed lines indicate clusters selected for reclustering and subsequent analysis. **B)** Marker genes used to identify clusters in (**A**). **C)** Violin plots of composite erythroid, myeloid and lymphoid signature scores in *gata2b+* HE cells from WT and caYAP embryos in GOF experiment (n = 39 WT, 37 caYAP cells, unpaired Student’s t-test *p ≤ 0.05). **D)** Violin plots of composite erythroid, myeloid and lymphoid signature scores in *gata2b+* HE cells from WT and yap1^−/−^ embryos in LOF experiment (n = 54 WT, 60 *yap1^−/−^* cells, unpaired Student’s t-test *p ≤ 0.05, ***p ≤ 0.001). **E)** qPCR on RNA from dissected tails for panel of hematopoietic transcription factors in embryos exposed to DMSO/1uM Yoda1 from 10ss–30hpf. Expression normalized to *18S* housekeeping gene (n = 5; full panel evaluated by two-way ANOVA, ***p<0.001; Error bars indicate SD). **F)** qPCR for human hematopoietic genes in human CD34+ cells exposed to DMSO or 0.5uM Yoda1 for 24hrs. Expression normalized to *18S* housekeeping gene (n = 4-6, unpaired Student’s t-test, *p ≤ 0.05).

**Supplemental Figure 5:**
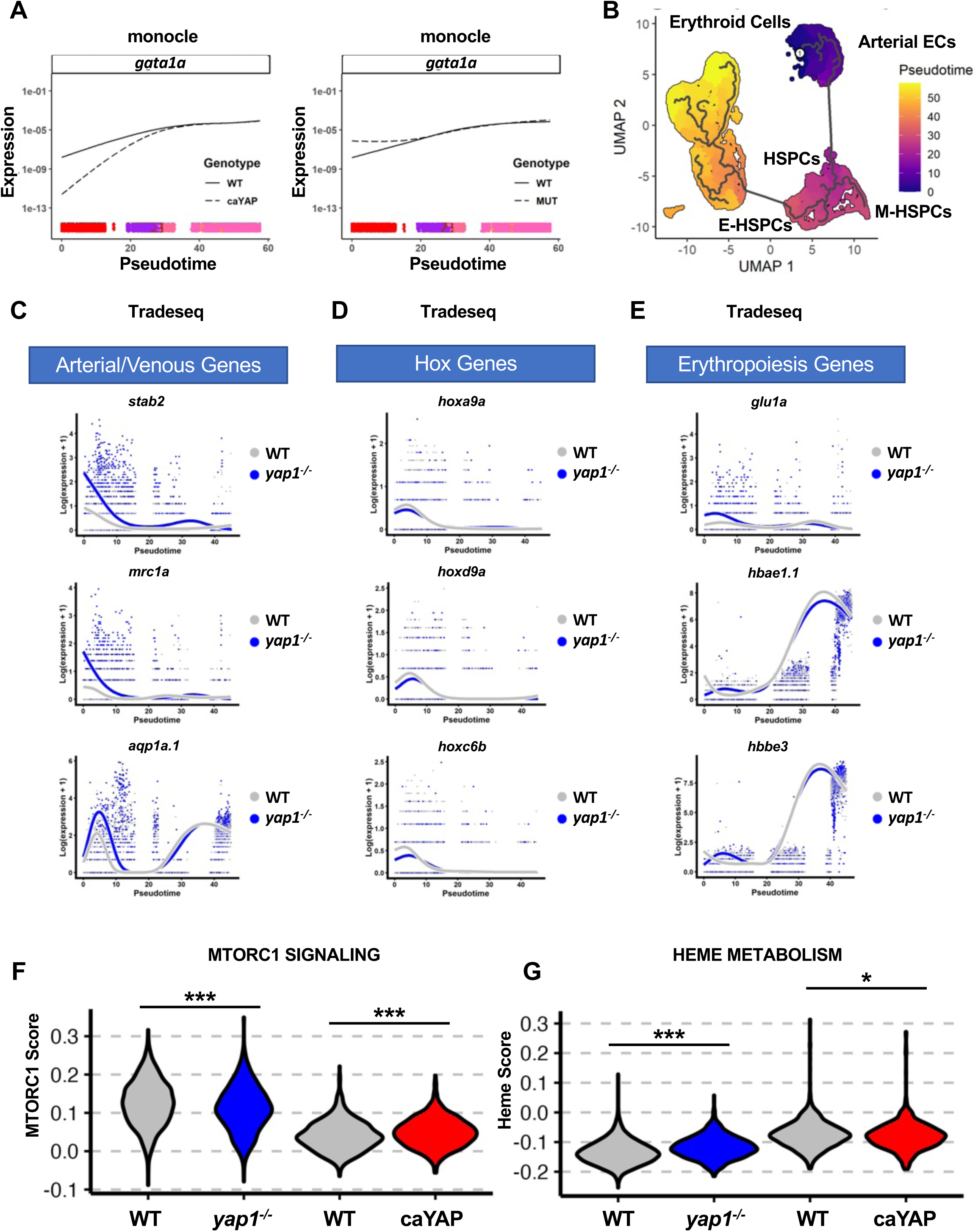
Loss of YAP causes EHT arrest by reversion to endothelial fates and blocked metabolic rewiring in HE cells. **A**) Erythrocyte master TF *gata1a* expression across pseudotime from Monocle in GOF and LOF experiments. **B)** Pseudotime trajectory from Monocle algorithm across the EHT UMAP from arterial to erythroid identity, passing through HSPC intermediates. **C)** Expression of arterial/venous genes across pseudotime from tradeseq in LOF experiments. **D)** Expression of HOX genes across pseudotime from tradeseq in LOF experiments. **E)** Expression of erythroid-associated genes across pseudotime from tradeseq in LOF experiments. **F)** Violin plots for composite scores of MTORC1 Signaling Hallmark pathway in the arterial endothelial cells from YAP GOF and LOF experiments (n = 1156 WT, 818 *yap1^−/−^* cells for LOF, 883 WT, 704 caYAP cells for GOF, unpaired Student’s t-test ***p ≤ 0.001). **G)** Violin plots for composite scores of Heme Metabolism Hallmark pathway in the arterial endothelial cells from YAP GOF and LOF experiments (n = 1156 WT, 818 *yap1^−/−^* cells for LOF, 883 WT, 704 caYAP cells for GOF, unpaired Student’s t-test *p ≤ 0.05, ***p ≤ 0.001).

**Supplemental Figure 6:**
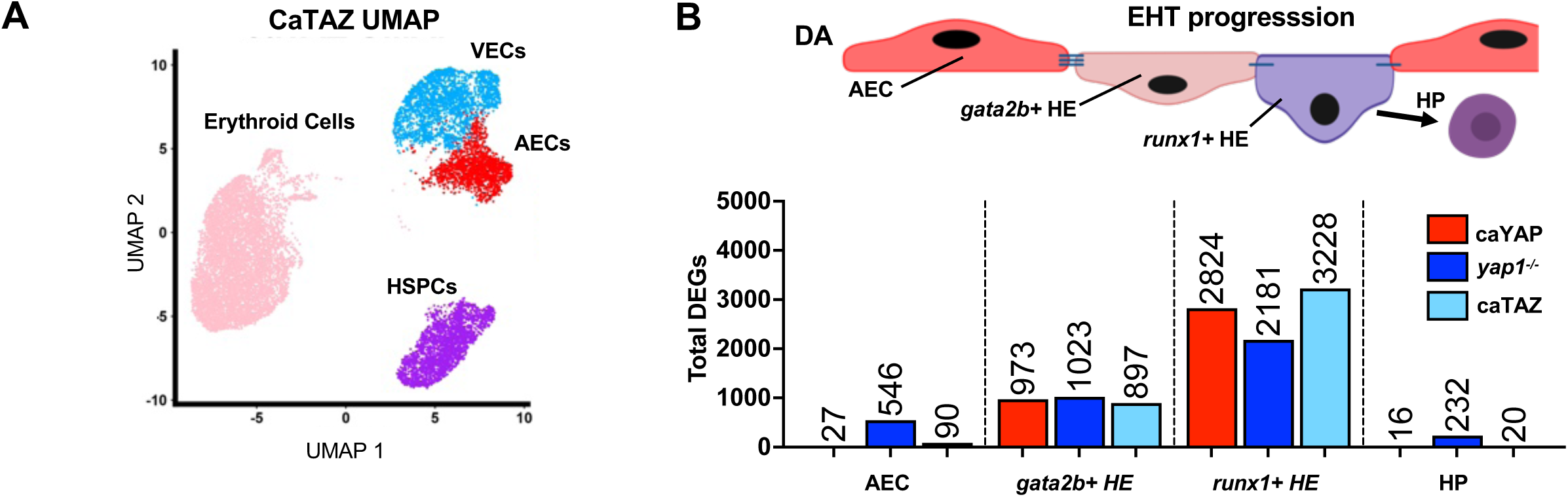
Sequencing of TAZ gain-of-function EHT cells shows selective effect of TAZ on augmenting endothelial signature. **A**) UMAP of reclustered Kdrl+ EHT and erythroid cell populations from WT and caTAZ GOF experiment. **B)** Graphical representation of EHT. The total arterial endothelial cell (AEC) cluster contains within it *gata2b*+ and *runx1+* HE cells, representing progressive states in the EHT continuum toward hematopoietic progenitors (HPs). Total DEGs in AEC, *gata2b+* HE, *runx1+* HE and HP cells are shown from FindMarkers() function with default parameters for caYAP, *yap1^−/−^* and caTAZ compared to respective WT controls.

**Supplemental Figure 7:**
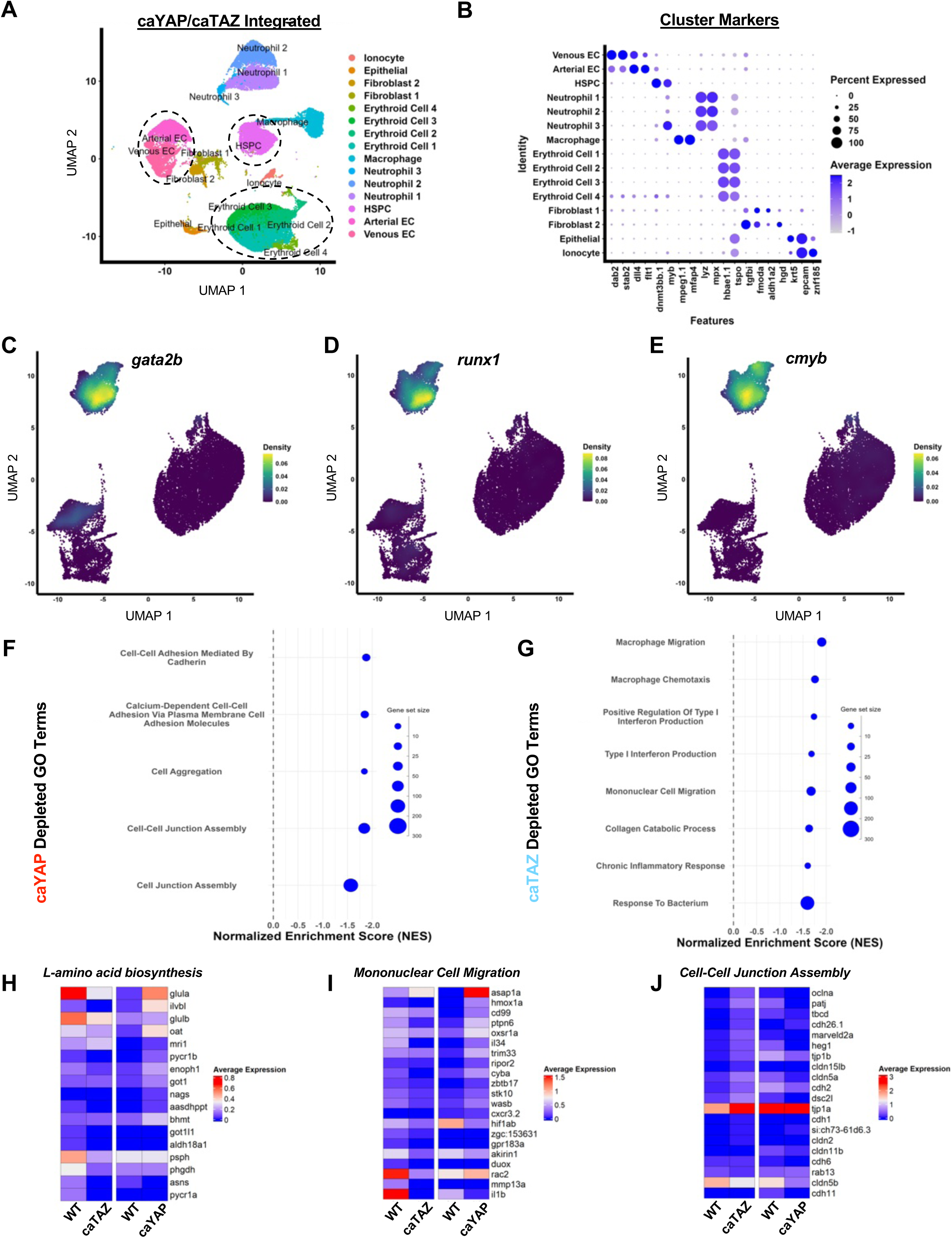
HSPC amplification by Piezo1 agonism is achieved by modulating shared YAP/TAZ proliferation pathways. **A**) UMAP following integration of the caYAP and caTAZ integrated datasets, circled populations were reclustered for use in figure 6. **B)** Cluster markers utilized to name clusters in A. **C-E)** Nebula plots displaying expression of *gata2b*, *runx1*, and *cmyb* across subsetted clusters **F)** GO:BP terms reflecting depleted adhesion programs in the caYAP dataset **G)** GO:BP terms reflecting depleted migration and immune response programs in the caTAZ dataset **H)** Genes within the “L-amino acid biosynthesis” GO:BP term plotted across caYAP and caTAZ datasets **I)** Genes within the “Monunuclear Cell Migration” GO:BP term plotted across caYAP and caTAZ datasets **J)** Genes within the “Cell-Cell Junction Assembly” GO:BP term plotted across caYAP and caTAZ datasets

**Supplemental Figure 8:**
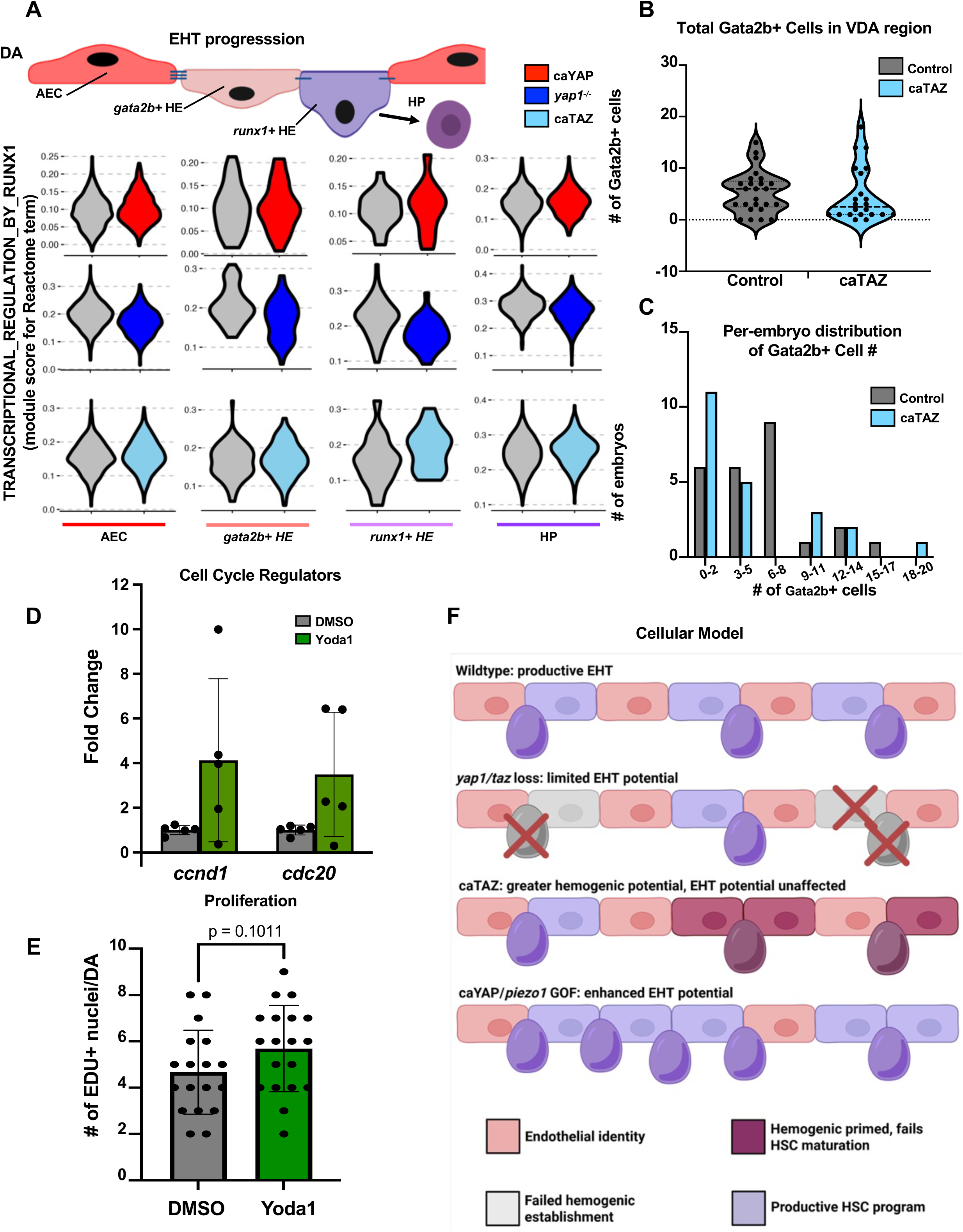
HSPC amplification by Piezo1 agonism is achieved by modulating shared YAP/TAZ proliferation pathways. **A**) Violin plots for ‘Transcriptional Regulation by Runx1’ Reactome term module score in AECs, *gata2b+* HE, *runx1+* HE and HP cells of different Hippo signaling genotypes (caYAP, *yap1^−/−^* and caTAZ) compared to WT. **B)** Quantification of Gata2b+ cells in confocal micrographs of control and caTAZ embryos at 48hpf. Dashed line indicates median cell number (N = 25 control, 23 caTAZ embryos). **C)** Histogram of number of embryos in bins of Gata2b+ cell numbers in the dorsal aorta (n = 25 control, 23 caTAZ embryos). **D)** qPCR on RNA from dissected tails for cell cycle genes *ccnd1* and *cdc20* in embryos exposed to DMSO/1uM Yoda1 from 10ss–30hpf. Expression normalized to *18S* housekeeping gene (n = 5; Error bars indicate SD). **E)** Quantification of EDU+ nuclei in the dorsal aorta from confocal images of WT *Tg(kdrl:EGFP)^s843^* embryos exposed to DMSO or 1uM Yoda1 from 10ss–32hpf. EDU pulse was delivered at 30hpf, embryos were fixed at 32hpf (n = 18, unpaired Student’s t-test; Error bars indicate SD). **F)** Cellular model for EHT dynamics in the dorsal aorta in WT and YAP/TAZ gain– and loss-of-function settings.

